# Single cell immunophenotyping identifies CD8^+^ GZMK^+^ IFNG^+^ T cells as a key immune population in cutaneous Lyme disease

**DOI:** 10.1101/2025.06.09.658661

**Authors:** Edel Aron, Hailong Meng, Paraskevas Filippidis, Alexia A. Belperron, Steven H. Kleinstein, Linda K. Bockenstedt

## Abstract

The skin lesion erythema migrans (EM) is the first clinical sign of Lyme disease, an infection due to the tick-transmitted bacterium *Borrelia burgdorferi* (*Bb*). Previously, we used scRNA-Seq to characterize the cutaneous immune response in the EM lesion, focusing on B cells. Here, with an expanded sample size, we profiled T cell responses in EM lesions compared to autologous uninvolved skin. In addition to CD4^+^ T cell subsets known to be abundant in the EM, we identified clonal expansion of CD8^+^ GZMK^+^ IFNG^+^ T cells that exhibited significant differential expression of interferon-regulated genes. This subset included IFNG^+^ cells with low cytotoxic gene expression, which may promote inflammation. While FOXP3^+^ regulatory T cells were also increased in EM, they exhibited little IL10 expression. In contrast, a CD4^+^ FOXP3^-^ tissue-resident T cell subset contained the largest population of cells with IL10 expression. Fibroblasts, endothelial cells, and pericytes were the principal cells that significantly differentially expressed key T cell-recruiting chemokines. These studies represent the first comprehensive interrogation of the cutaneous T cell response to *Bb* infection using single cell transcriptomics with adaptive immune receptor sequencing, providing insight into the skin barrier defense and orchestration of the immune response to this vector-borne pathogen.

## Introduction

Lyme disease (LD), caused by infection with *Ixodes* tick-transmitted spirochetes of the *Borrelia burgdorferi sensu lato (Bb)* family (1), is the most common vector-borne disease in North America and Europe (2). An expanding erythematous skin lesion known as erythema migrans (EM) is the first clinical sign of infection in 70-80% of patients and arises at the bite site within one month of infection. If unchecked at this barrier site of entry, *Bb* may disseminate to cause neurologic, cardiac, and rheumatologic manifestations. The infection readily responds to antibiotics, although some people experience protracted symptoms, and a minority develop post-infectious sequelae. These include persistence of inflammatory arthritis despite antibiotics (post-infectious Lyme arthritis) with autoimmune features that resolve with anti-rheumatic therapies (3–5) and a post-treatment LD syndrome of cognitive dysfunction, musculoskeletal pain, and fatigue of unclear etiology (6–8). There is strong evidence for Type II interferon (IFN) responses contributing to disease pathogenesis from the earliest stages when EM is present (9–11), as well as for dysregulation of such responses leading to post-infectious Lyme arthritis (12). Understanding the orchestration of host defense at the skin barrier site is a key factor in preventing LD and its adverse sequelae.

EM appears at a time when the actions of tick saliva on local skin immunity are estimated to wane (approximately 2 weeks after tick detachment) (13). At this stage, histopathology and flow cytometry studies of the lesion reveal that IFNγ-producing T cells, monocytes/macrophages, and DCs comprise the main cells infiltrating dermal tissues, along with a variable presence of B cells and plasma cells (14–16). A transcriptomic microarray and pathway analysis revealed the dominance of an IFN signature, with gene expression of IFNG but not IFNA detected in the skin (9). B cells are a critical host defense against *Bb* infection yet are significantly underrepresented in normal skin in comparison to other cell types (17, 18). To further investigate the role of B cells in host defense at this barrier site, our group performed single cell transcriptomics (scRNA-Seq) alongside B cell receptor (BCR) and T cell receptor (TCR) sequencing to deeply interrogate the cutaneous immune response in LD, with an emphasis on the B cell response (19). When comparing the composition of cells within EM lesions to those in autologous uninvolved skin, we found that the lesions were enriched for B cells along with plasma cells. These B cells had clonal relatives in the blood, differentially expressed MHC class II genes, and exhibited a significant signature for MHC class II antigen presentation and IFNG signaling. The presence of a large unmutated IgM memory cell population in the EM lesions supported an extrafollicular B cell response that may be critical for controlling spirochetes at this early stage of infection. A limited analysis of EM T cells revealed clones that could be traced to the circulation. These cells displayed gene signatures of cell trafficking and co-stimulation with CD28 as well as antigen-driven selection, as evidenced by clonal expansion. Given the prominence of T cells in the EM lesions, in this study we sought to conduct a deeper interrogation of the cutaneous T cell response in LD using an expanded sample size. Our results demonstrate involvement of a variety of T cell subsets, including a novel contribution of CD8^+^ GZMK^+^ IFNG^+^ T cells, and through cytokine and chemokine analyses, provide new insights into the role skin cells play in orchestrating host defense against this vector-borne pathogen.

## Results

### All defined T cell subsets are enriched in the EM lesion

To characterize the cellular composition of the EM lesion, we used the 10x Genomics platform to perform scRNA-Seq along with adaptive immune receptor sequencing (20) on cells from whole skin digests of EM and uninvolved autologous (control) skin biopsies from 13 participants with acute LD (Table 1) (20). Analysis of the combined data from EM lesions and control skin samples for all participants identified 42 cell clusters (Figure 1A, Supplemental Figure 1A). These clusters corresponded to 13 distinct cell types based on the expression of canonical marker genes (Figure 1B, Supplemental Figure 1B): B cells, cycling dendritic cells, endothelial cells, fibroblasts, keratinocytes, Langerhans cells, macrophages, mast cells, melanocytes, natural killer (NK) cells, nerve cells, pericytes, and T cells.

**Table 1.** Characteristics of Study Subject Participants.

| Dataset | Participant Number | Age (years) | Sex | EM Lesion(s) | Lesion Size (cm) | Lesion Site(s) | Paired Control |
| --- | --- | --- | --- | --- | --- | --- | --- |
| 1 | 192561 | 68 | F | Single | 9 | Left popliteal fossa | Yes |
| 1 | 192563 | 63 | F | Single | 4 | Right proximal triceps | No |
| 1 | 192564 | 68 | F | Single | 6.5 | Left axilla | Yes |
| 1 | 192565 | 70 | F | Single | 5 | Right popliteal fossa | Yes |
| 1 | 192566 | 40 | M | Multiple | 10<sup>A</sup> | Left upper back, anterior abdomen | Yes |
| 1 | 192567 | 72 | F | Single | 10 | Right ankle | Yes |
| 2 | 202936 | 31 | M | Single | 7 | Left flank | Yes |
| 2 | 202937 | 61 | M | Single | 20 | Right abdomen | Yes |
| 2 | 202938 | 65 | F | Single | 4 | Right thigh | Yes |
| 3 | 202939 | 72 | F | Single | 9 | Left axilla | No |
| 3 | 202940 | 64 | F | Single | 15 | Right shoulder | Yes |
| 3 | 202941 | 70 | F | Single | 12 | Left back | Yes |
| 3 | 202943 | 76 | M | Single | 20 | Left thighs | Yes |

<sup>A</sup> The recorded size reflects the largest lesion.

**Figure 1.**
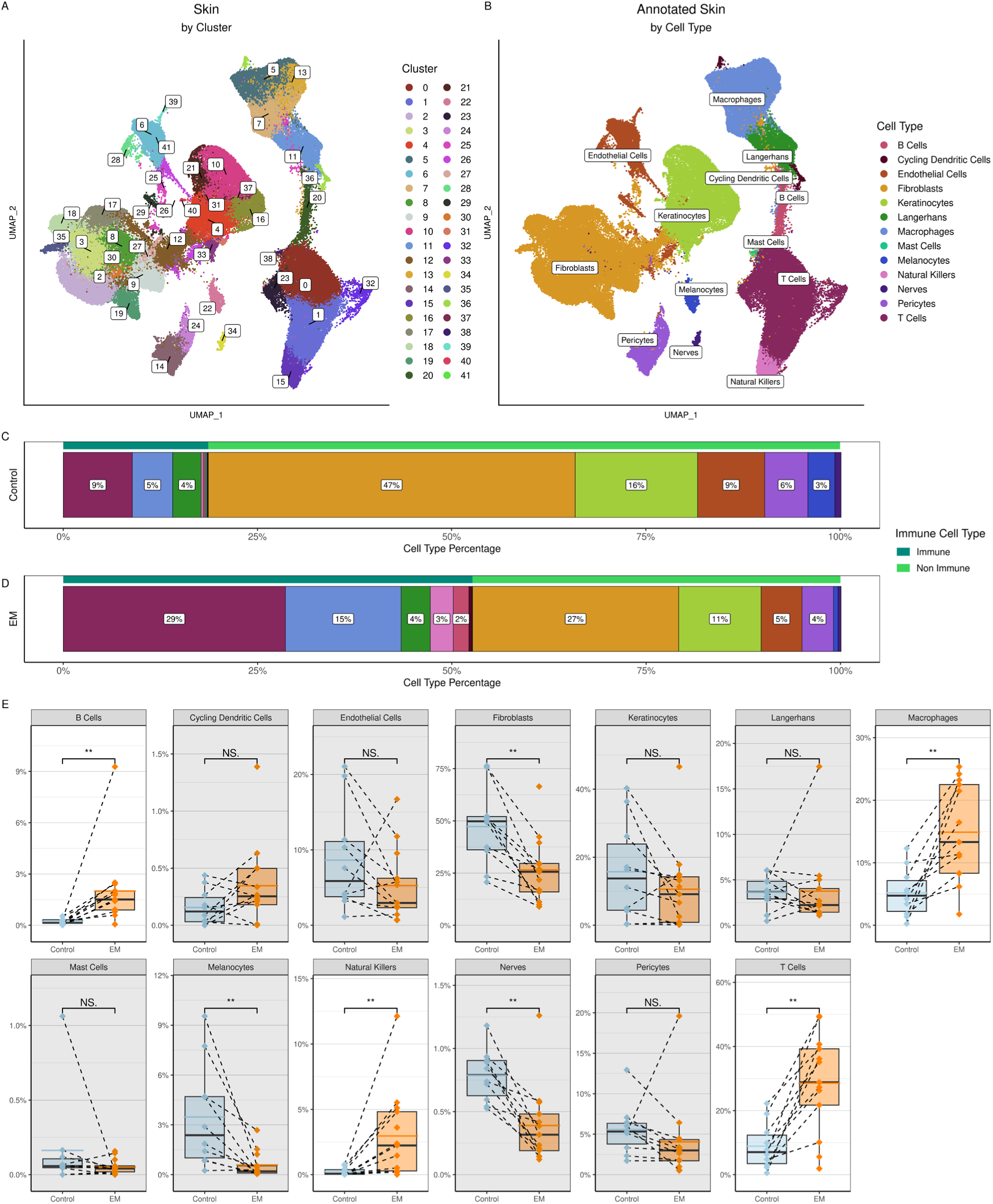
Clustering and annotation of the skin cells. **A)** UMAP representing clustering of the combined control and EM skin data. At a resolution of 1.0, there were 42 clusters formed. **B)** UMAP of skin cell types annotated using canonical marker genes. **C)** and **D)** Proportions of cell types within the control and EM samples, respectively, grouped by immune (dark green bars) and non-immune cell types (light green bars) then arranged in decreasing order by percentage. Proportions were averaged by participant for each cell type. Colors correspond to the cell types in B). **E)** Comparison of the frequencies of each cell type between sample types (control vs. EM). Points represent results from individual participants and the dashed lines connect paired samples. The solid black horizontal lines within each box plot indicate the median and the colored (blue or orange) lines indicate the mean for each sample type and cell type across participants. Gray rectangles indicate cell types that did not have significant increases in EM compared to control samples. Significances were calculated using a paired Wilcoxon signed-rank test (***P < 0.001; **P < 0.01; *P < 0.05).

As expected, the uninvolved skin samples were mainly composed of fibroblasts (47%) and keratinocytes (15%) (Figure 1C) (21). T cells (10%) dominated the immune cell population, with B cells (<1%) rarely detected, in line with previous studies (18, 19). The proportions of most immune cells in the EM lesion increased in comparison to uninvolved skin, with T cells remaining the largest cell population overall (29%) (Figure 1C, D). Significant increases in the frequencies of individual immune cell types were found in the EM. Among those increased in the EM lesion were the B cells, macrophages, NK cells, and T cells (Wilcoxon signed-rank test, p < 0.002), with corresponding decreases in frequencies of non-immune cells (Figure 1E). As a result of these changes, more than half (56%) of the cells in the EM lesions were immune-related, as opposed to 21% in the control skin.

T cells were further divided into 26 clusters (Figure 2A) and annotated using marker genes into 11 distinct subtypes (Figure 2B, Supplemental Figure 2). While we were able to assign clear labels to nine of these cell types, two subsets of CD4^+^ and CD8^+^ T cells were labeled as “undefined” due to the lack of additional gene markers or cell surface protein data that would permit further characterization. In addition to CD4^+^, CD8^+^, and FOXP3^+^ regulatory T cell subsets, the nine identified T cell subsets included double negative (CD4^-^ CD8^-^) T cells, dividing T cells, naïve T cells, and a group of cells that upregulated heat shock protein (HSP) genes, designated as HSP^hi^ T cells (Figure 2B). Although the HSP^hi^ T cells may be stressed or dying cells that are often seen in scRNA-Seq data due to tissue processing (22), these cells also may represent a population experiencing cellular stressors in the context of skin infection. No NKT cells were identified. Even though the proportions of the T cell subtypes varied between the EM and control skin cells (Figure 2C, D), all defined T cell subsets were significantly increased in the EM when compared to controls (Figure 2E).

**Figure 2.**
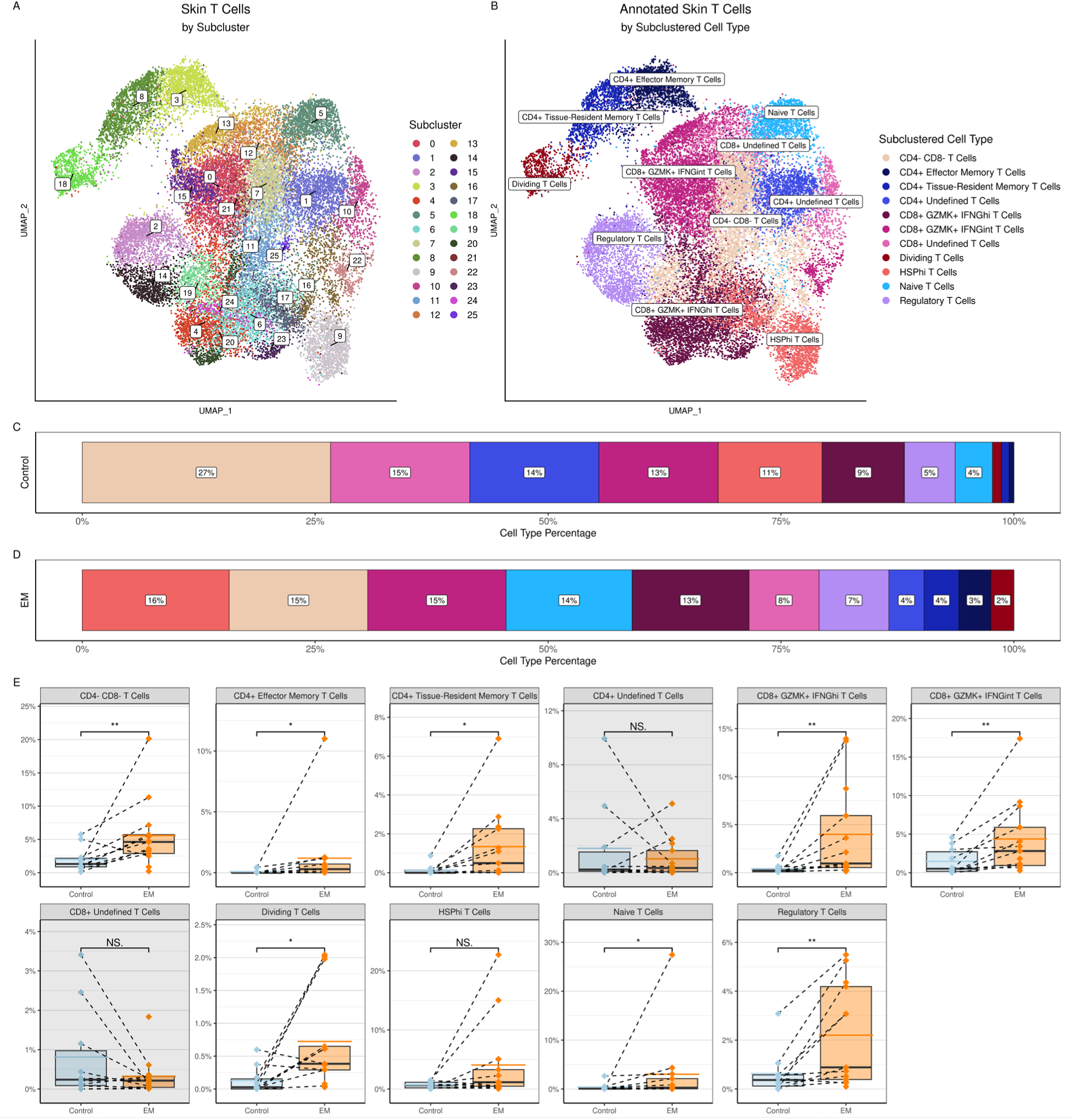
Clustering and annotation of the skin T cells. **A)** UMAP representing clustering of the subclustered T cells from the combined control and EM skin data; at a resolution of 1.5, there were 26 clusters formed. **B)** UMAP of skin T cell types annotated using canonical marker genes. **C)** and **D)** Proportions of cell types within the control and EM samples, respectively, arranged in decreasing order by percentage. Proportions were averaged by participant for each cell type. Colors correspond to the cell types in B). **E)** Comparison of the frequencies of each cell type between sample types (control vs. EM). Cell counts are relative to all of the skin cells, not just the total T cells. Points represent results from individual participants and the dashed lines connect paired samples. The solid black horizontal lines within each box plot indicate the median and the colored (blue or orange) lines indicate the mean for each sample type and cell type across participants. Gray rectangles indicate the cell types that did not have significant increases in EM compared to control samples. Significances were calculated using a paired Wilcoxon signed-rank test (***P < 0.001; **P < 0.01; *P < 0.05).

CD4^+^ T cells were subdivided into two groups representing CD4^+^ effector memory and CD4^+^ tissue-resident memory subsets, in addition to the “undefined” CD4^+^ T cell subset. The distinction between CD4^+^ T effector memory cells and the CD4^+^ tissue-resident memory subsets was primarily based upon expression of integrins and cytotoxic markers, as other distinguishing markers such as CD69 were not detected in either subset. Characterization of the effector memory T cell subset revealed the presence of the tissue-residency marker CXCR6 and the memory T cell marker KLRB1/CD161. This subset also expressed the exhaustion marker PDCD1 and the post-activation inhibitory molecule CTLA4, but not cytotoxic markers such as GZMA or tissue-homing markers such as CCR7. The CD4^+^ tissue-resident memory T cells also expressed CXCR6, KLRB1, and PDCD1, but could be distinguished from the effector memory T cells by their high expression of the signaling regulator RGS1 and integrins such as ITGAE/CD103 and ITGA4 (as well as low expression of CCR7), suggesting an activated or cytotoxic phenotype. This subset also included activated T cells with the potential for Th1-like function as implied by expression of GZMA, IFNG, and the transcription factor TBX21/T-bet.

We initially defined Tregs by their expression of FOXP3. FOXP3^+^ Tregs have been identified in peripheral blood and synovial fluid of people with Lyme arthritis, and a reduction in their numbers and ability to suppress IFNG production has been associated with immune dysregulation leading to post-infectious Lyme arthritis (23). To date, there have been no direct assessments of Treg cells in EM lesions, although their presence has been inferred by the detection of IL10 mRNA in EM immune cells by in situ hybridization and IL-10 in blister fluid from EM lesions (14, 15). We therefore characterized T cell subsets using differential gene expression of IL10 in addition to IFNG, as FOXP3 ^+^ T cells can upregulate IFNG in the context of Th1-mediated immune responses and still retain their immunosuppressive potential (24, 25).

Although multiple T cell subsets exhibited expression of IL10, FOXP3^+^ Tregs (which were not exclusively confined to the CD4^+^ subset) were not the main source of IL10 as expected based on previous studies (23). Instead, the CD4^+^ tissue-resident memory T cells, which were FOXP3^-^, had the highest proportion of cells expressing IL10 among all T cell types (Supplemental Figure 3A-F). These CD4^+^ FOXP3^-^ IL10^+^ tissue-resident memory T cells are reminiscent of Type I regulatory (Tr1) T cells as they included cells that express CTLA4, LAG3, and TGFB (as well as ICOS, PDCD1, TGFB, and TOX2), but not BCL6, a transcription factor involved in germinal center formation (26–28). Tr1 T cells can regulate immune responses through gene expression as well as through induction of apoptosis (29–31). Accordingly, we found that TNF-related apoptosis-inducing ligand TNFSF10/TRAIL was expressed within this population, suggesting that these T cells may suppress inflammation through apoptosis in addition to the secretion of IL10 (Supplemental Figure 3G-I).

IFNG was detected in multiple CD4^+^ T cell subsets, especially CD4^+^ tissue-resident memory cells as predicated by their activation markers and expression of TBX21/T-bet (Supplemental Figure 3J-L). We also detected IFNG expression in FOXP3^+^ Tregs. However, we unexpectedly found the greatest proportion of IFNG expression within the CD8^+^ GZMK^+^ T cell population. This cluster could be subdivided based on the level of IFNG expression into two groups, namely CD8^+^ GZMK^+^ IFNG^hi^ T cells and CD8^+^ GZMK^+^ IFNG^int^ T cells. CD8^+^ T cells have been identified in EM lesions in prior studies and are considered to serve cytotoxic functions, although they have not been fully characterized (15, 32). To assess the cytotoxic potential of these cells, we analyzed the expression of PRF1 (perforin) and GZMB in each of these subclusters and found that while these genes were expressed by cells within both populations, we also observed a sizeable number of cells with very low or nondetectable levels of these cytotoxic genes. The CD8^+^ GZMK^+^ IFNG^hi^ group had a larger proportion of these GZMB^low^ PRF1^neg^ cells in comparison to the CD8^+^ GZMK^+^ IFNG^int^ T cells (Supplemental Figure 4A-D). These CD8^+^ GZMK^+^ IFNG^hi^, GZMB^low^, PRF1^neg^ cells were enriched in the EM lesion and are reminiscent of tissue-enriched expressed GZMK^+^ T cells (T_teK_ cells) that were initially identified in rheumatoid synovium and subsequently found at barrier sites such as the lung, intestine, and skin (33).

The presence of cells expressing PRF1 and GZMB within CD8^+^ GZMK^+^ IFNG^hi^ and CD8^+^ GZMK^+^ IFNG^int^ subclusters suggested that these populations could also contain terminal effector memory re-expressing CD45RA (TEM_RA_) cells, which have high cytotoxic potential (37). As we had no corresponding protein-based phenotyping data, we computationally inferred the presence of spliced PTPRC/CD45 variants to identify a possible TEM_RA_ subset (see Methods). We found that a large proportion of possible TEM_RA_ cells were in the CD8^+^ GZMK^+^ IFNG^int^ subset (Supplemental Figure 5). Interestingly, these T cells also contained a higher proportion of cells expressing the CD45RB isoform, which has been associated with virus-specific CD8^+^ memory T cells (38).

### CD8^+^ GZMK^+^ IFNG^hi^ T cells exhibit differential expression of genes distinct from CD8^+^ GZMK^+^ IFNG^int^ T cells in the EM lesion

CD8^+^ GZMK^+^ IFNG^hi^ T cells may represent cells with similar differentiation trajectories as CD8^+^ GZMK^+^ IFNG^int^ T cells in the EM lesion. Because trajectory analysis was inconclusive, we ran a pseudobulk differential gene expression analysis comparing these two cell populations within the EM lesion to gain insight into which genes distinguish CD8^+^ GZMK^+^ IFNG^hi^ T cells from the CD8^+^ GZMK^+^ IFNG^int^ T cells other than the relative amount of IFNG expression (Figure 3). In addition to the IFNG, CD8^+^ GZMK^+^ IFNG^hi^ T cells upregulated several HSP genes, including members of the HSP70 (HSPA6, HSPA1B, HSPA1A) and the HSP70-partner HSP40 (DNAJB1, DNAJB4) families, suggesting that they were exhibiting cell stress or even potentially inducing cytotoxicity (39–41). This population also included genes that promote (CCL3, CCL4, CD83) or modulate (NFKBIE, EGR1, TRAF3IP3, NKRF) inflammatory responses or affect T cell migration (RGS16, RGS2) (42–45). In contrast, CD8^+^ GZMK^+^ IFNG^int^ T cells upregulated genes involved with T cell activation and survival (TNFRSF9/4-1BB), cytotoxicity (PRF1), tissue localization (CXCR6), and regulation (PDE4A, TGFB1) (46). Thus, these cells may represent populations that evolve from common precursors yet achieve different functions depending on the skin microenvironment.

**Figure 3.**
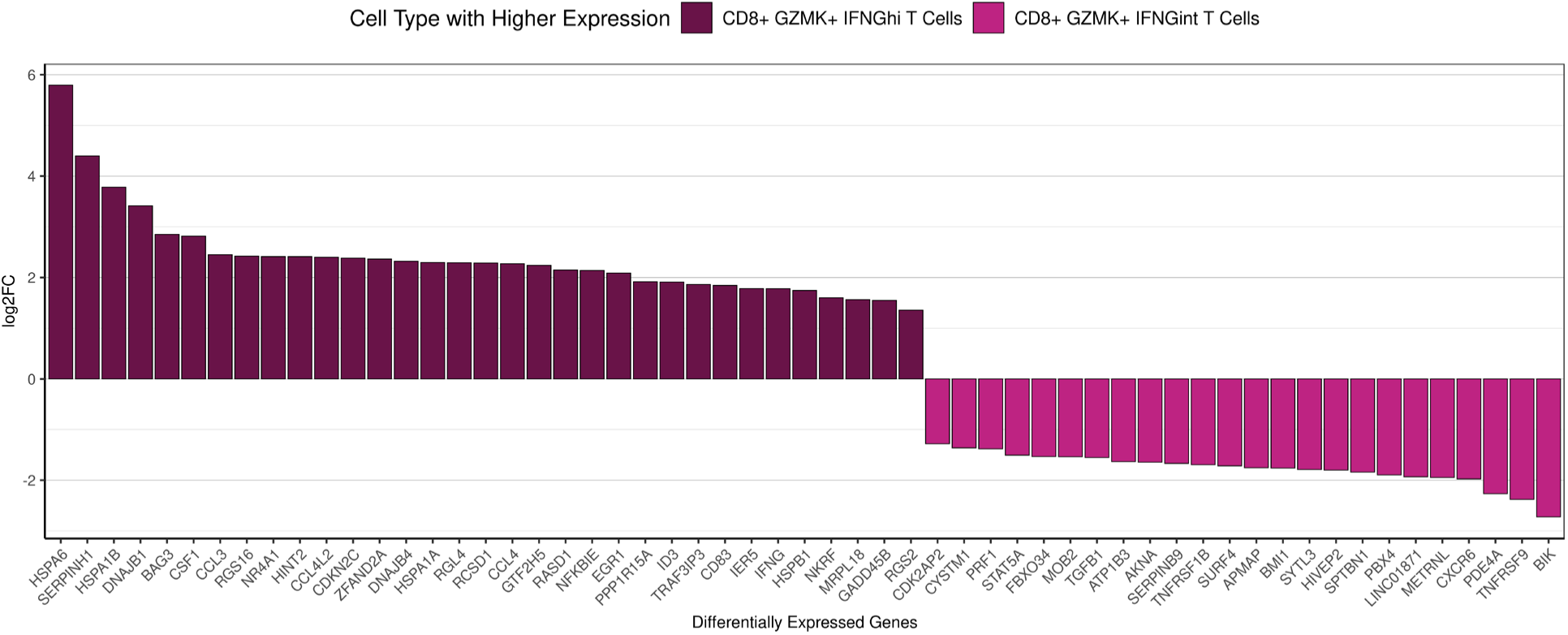
Differentially expressed genes between CD8^+^ GZMK^+^ IFNG^hi^ T cells and CD8^+^ GZMK^+^ IFNG^int^ T cells in the EM lesion only. Bar plot showing the top differentially expressed genes in EM compartments of the CD8^+^ GZMK^+^ IFNG^hi^ T cells and CD8^+^ GZMK^+^ IFNG^int^ T cells with an adjusted p < 0.05, arranged by log2FC. Note that a negative log2FC reflects genes being expressed in the cell type being compared to, not downregulation.

### EM T cells are clonally expanded, and a subset is shared with the blood

Immune repertoire analysis revealed that the increased T cell count in the EM as compared to the control skin results from both clonal expansion as well as an increase in the number of clonotypes (Supplemental Figure 6). To better understand this expansion and investigate potential migration between the skin and the blood, we quantified unique clonotypes across tissues. As expected, we observed that the blood T cells had the greatest number of clonotypes but 96.5% of them were singletons (not expanded) (Figure 4, Supplemental Figure 7). Only 1.99% of these T cell clonotypes were shared with the EM lesion and 0.28% with the control skin (Figure 4A). A two-sample proportion test revealed that the overlap between the blood and the EM clonotypes was significantly more than between the blood and the control skin clonotypes (p = 0.004).

**Figure 4.**
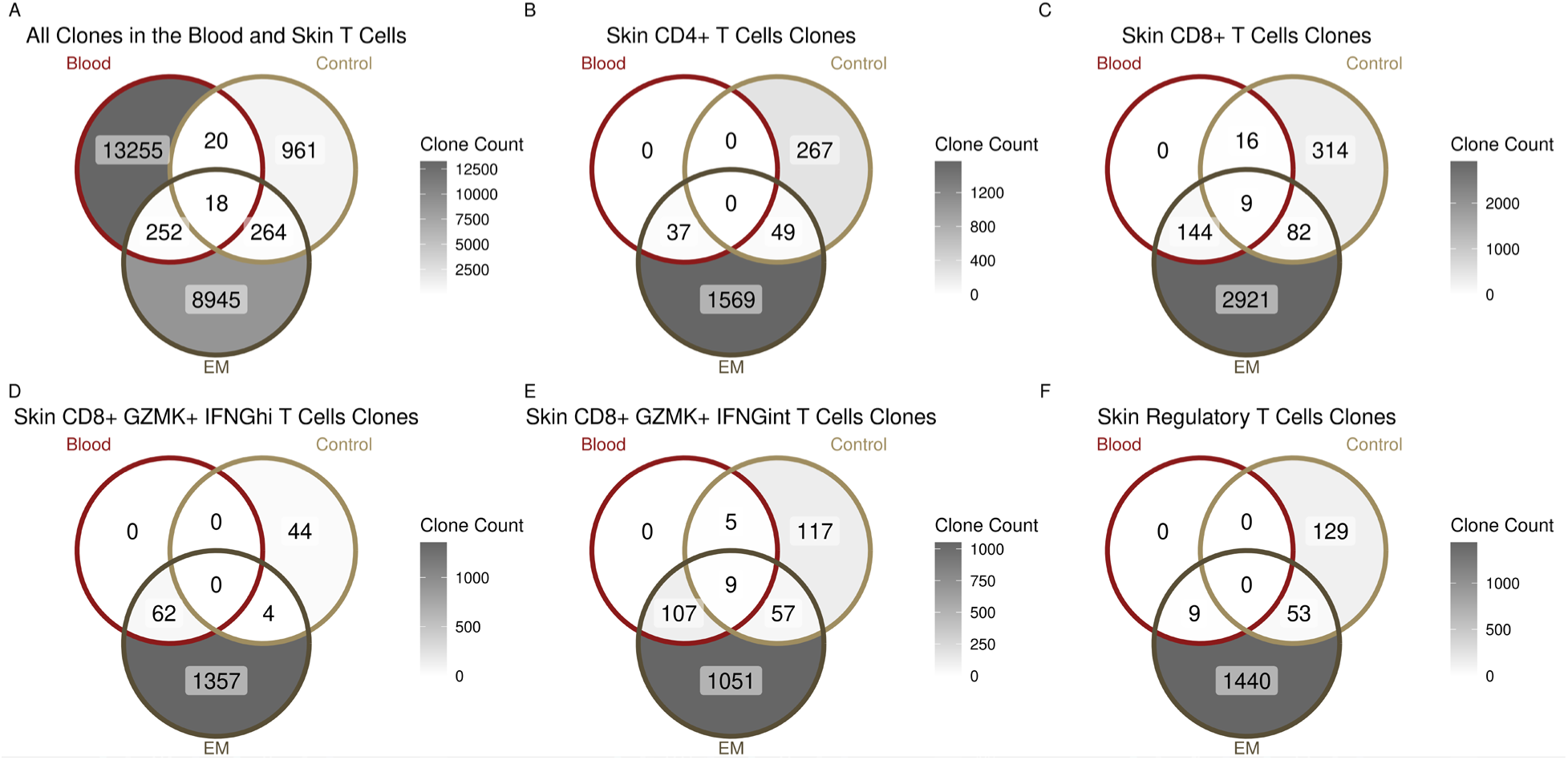
T cell clonal overlaps between the blood and the skin. **A)** Clonal overlaps between all of the blood T cells (dark red) and the skin T cells, with the latter split into EM (dark brown) and control skin (light brown). **B)** and **C)** Clonal overlaps between the blood T cells and the skin CD4^+^ T cells (CD4^+^ effector memory T cells, CD4^+^ tissue-resident memory T cells, and CD4^+^ undefined T cells) or CD8^+^T cells (CD8^+^ GZMK^+^ IFNG^hi^ T cells, CD8^+^ GZMK^+^ IFNG^int^ T cells and CD8^+^ undefined T cells) respectively. **D), E),** and **F)** Clonal overlaps between the blood T cells and the skin CD8^+^ GZMK^+^ IFNG^hi^ T cells, CD8^+^ GZMK^+^ IFNG^int^ T cells, and Tregs, respectively.

We next determined which T cell subsets within the EM lesion had clonotypes shared with the blood and distinct from those in control skin. A minority of the total EM CD4^+^ T cell clonotypes (2.24%) and none of the control skin clonotypes were traced to the circulation (p = 0.014) (Figure 4B), in comparison to 5.94% of EM and 4.85% of control skin CD8^+^ T cell clonotypes (p = 0.017) (Figure 4C). When comparing CD8^+^ GZMK^+^ cells, a greater percentage of EM CD8^+^ GZMK^+^ IFNG^int^ T cell clonotypes (9.48%) than EM CD8^+^ GZMK^+^ IFNG^hi^ T cell clonotypes (4.36%) shared TCRs with cells in the blood (Figure 4D, E). Moreover, both EM and control skin CD8^+^ GZMK^+^ IFNG^int^ T cell clonotypes were detected in the blood (Figure 4E), with the overlap between the blood and the EM being significantly greater than the blood and the control (p = 0.048). In contrast, none of the control skin CD8^+^ GZMK^+^ IFNG^hi^ T cell clonotypes were detected in the blood. We also examined the FOXP3^+^ Tregs subset and found only EM Treg clonotypes in the blood, similar to the pattern seen with CD4^+^ and CD8^+^ GZMK^+^ IFNG^hi^ T cell clonotypes (Figure 4F). These findings indicate that CD8^+^ GZMK^+^ IFNG^int^ clonotypes include cells that exhibit greater migratory capacity between skin and blood in comparison to other T cell subsets examined.

### Non-immune cells express T cell-recruiting chemokines in EM lesions

We next examined which skin cell types might be responsible for recruiting the T cells to the EM. Prior studies reported that DCs and macrophages were the principal immune cells attracting T cells to sites of *Bb* infection in tissues (13, 47). However, dermal fibroblasts are also important sources of lymphocyte-attracting cytokines and chemokines, as has been demonstrated in studies of autoimmune and inflammatory skin diseases (48–50). Thus, we assessed chemokine expression in both immune and non-immune cell subsets to identify cell types that may attract T cells and other immune cells to the EM lesion. We used differential gene expression analysis to compare the expression of all chemokine ligand and receptor genes in each cell type in the EM and the control skin (Figure 5, Supplemental Figure 8). We found significant upregulation of genes in non-immune cells (endothelial cells, fibroblasts, and pericytes) as well as immune cells (B cells, CD8^+^ GZMK^+^ IFNG^hi^ T cells, macrophages, NK cells). In contrast, Langerhans cells and macrophages were the only cells exhibiting significant downregulation of chemokines, predominantly those that recruit neutrophils (CXCL1, CXCL2, CXCL3, CXCL5, and CXCL8), consistent with the paucity of neutrophils found on EM histopathology (51).

**Figure 5.**
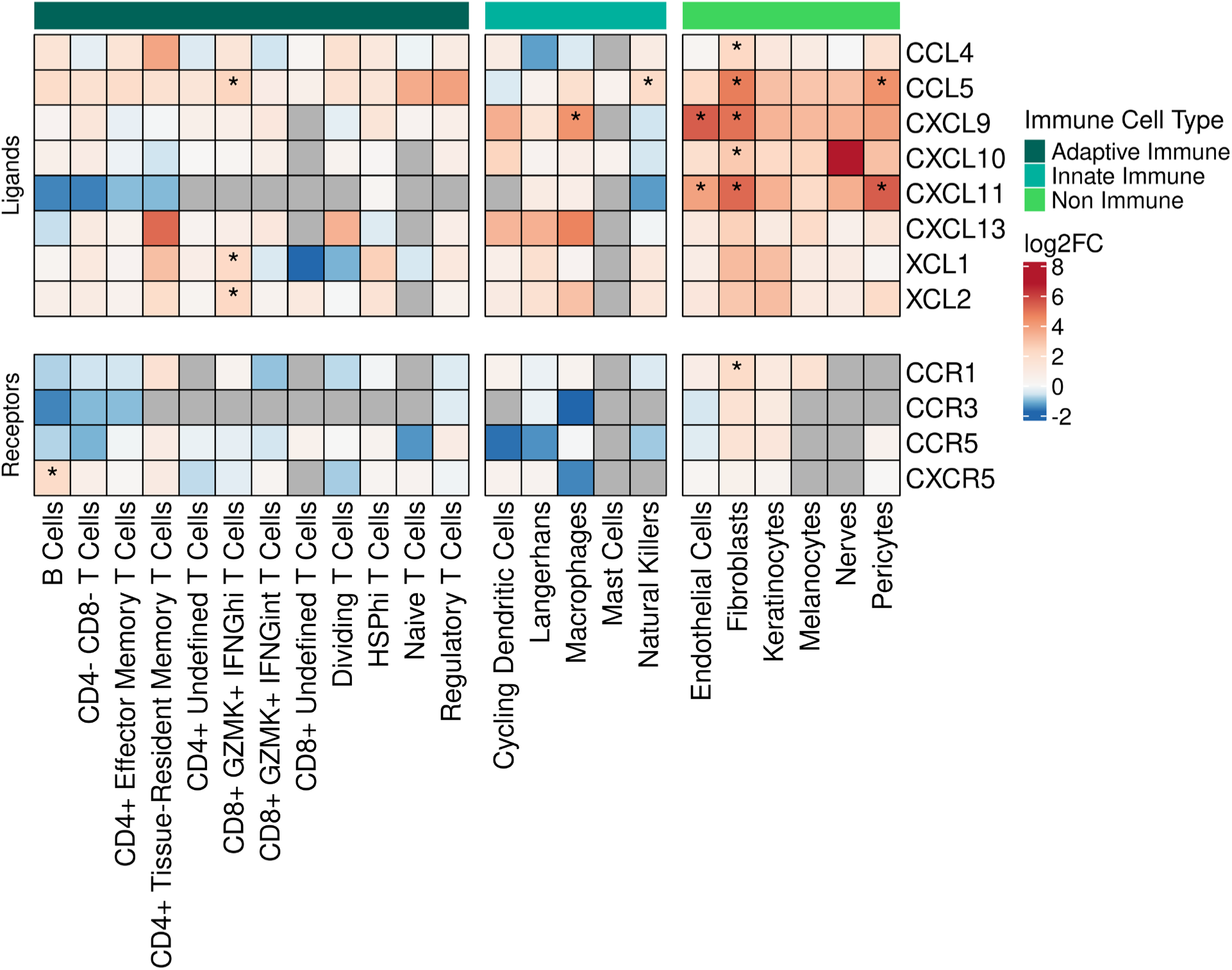
Gene expression of chemokine ligand and receptor genes in the skin. Heatmap showing the average fold change of the expression of select chemokine-related genes across the full set of skin cell types. The bar on the top indicates whether or not the cell types being plotted are immune-related (dark green for adaptive immune cell types, cyan for innate immune cell types, and bright green for non-immune cell types). Within the heatmap itself, red indicates upregulation, and blue indicates downregulation, with asterisks denoting statistically significant changes (FDR < 0.05). Gray cells indicate no expression of the relevant gene for that cell type (not applicable). Differential expression was calculated between the EM and control portions of each cell type.

Fibroblasts significantly upregulated the greatest number of chemokine genes, including the T cell-recruiting chemokine CCL5 (52) and its receptor CCR1, lymphocyte chemoattractants CXCL9, CXCL10, and CXCL11, as well as pro-inflammatory macrophage attractants CCL4, CCL18, CCL19, CCL26, and the neutrophil attractant CX3CL1. Significantly upregulated lymphocyte recruiting chemokine genes were also found in endothelial cells (CXCL9 and CXCL11) and pericytes (CXCL11), but macrophages only upregulated CXCL9. The only significantly differentially expressed chemokine genes by T cells were found in the CD8^+^ GZMK^+^ IFNG^hi^ subset and included XCL1 and XCL2, which recruit conventional DCs important for cross-presentation of antigen (53). In addition, this T cell subset significantly upregulated the T cell chemoattractant CCL5. Overall, several chemokine ligand and receptor genes were significantly expressed across a variety of cell types, with many non-immune cell types differentially expressing at least one gene (Supplemental Figure 8). However, most chemokines specifically involved with T cell recruitment were found to be upregulated in non-immune cells.

### Interferon response pathways are enriched in the EM lesion, but only IFNG expression is upregulated

In a previous study, interferon signaling (including the upstream regulators IFNA and IFNG) was implicated by pathway analysis of bulk EM transcriptomes, with only IFNG expression detected in the lesion (9). Since many of the chemokines we observed to be upregulated in the EM lesion are induced by interferons, we sought to gain a better understanding of the biological pathways involved by performing gene set enrichment analysis (GSEA) comparing EM and control skin across cell types using Human Molecular Signatures Database’s Hallmark reference. GSEA showed a total of 43 immune pathways with positive enrichment in at least one cell type (out of 50 total Hallmark pathways), including those involving apoptosis, complement, interleukin, JAK/STAT signaling, IFN responses, TGFB signaling, and TNFA signaling (Figure 6). Both the IFNA and the IFNG response pathways were significantly enriched in almost all cell types, as was the TNFA signaling via the NFKb pathway (except in the cycling DCs and the macrophage populations). When examining cells impacted by IFN signaling, we found that several individual interferon-regulated genes (IRGs) were significantly upregulated within the CD8^+^ GZMK^+^ IFNG^hi^ T cells, including IFITM1, IFITM2, IRF1, IRF7, ISG20, and STAT1 (Figure 7). Most of these genes were also significantly upregulated within B cells, along with genes associated with IFN signaling and downstream effects of IFN such as JAK1, NFKB, and TNF.

**Figure 6.**
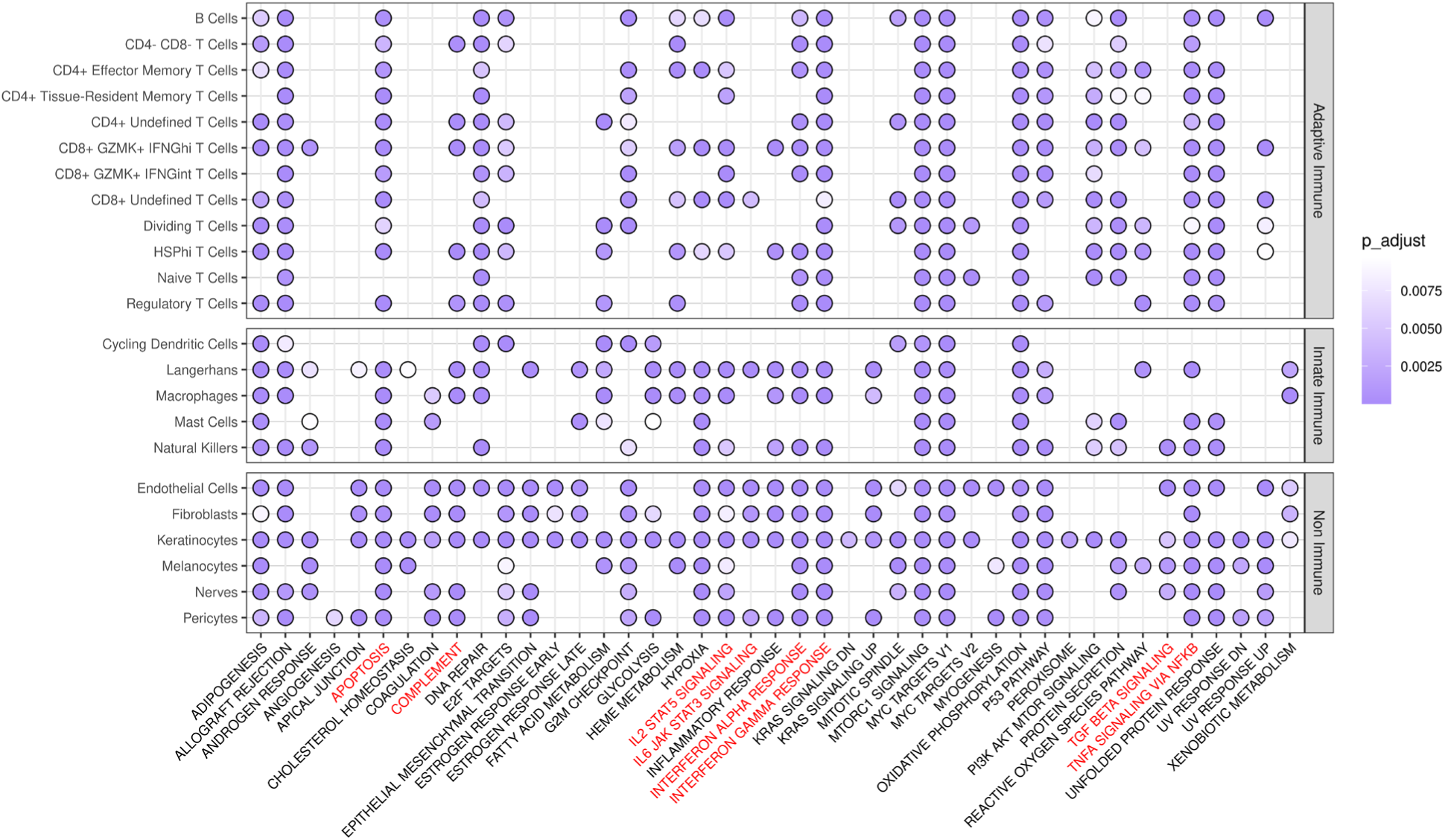
GSEA significance analysis of using the MSigDB Hallmark database in the skin. Gene set enrichment analysis (GSEA) was performed using the Hallmark database from the Human Molecular Signatures Database (MSigDB). Purple circles indicate significance (Bonferroni-corrected p-value < 0.01) and immune pathways of note are highlighted in red. Only pathways with significance in at least one cell type are shown.

**Figure 7.**
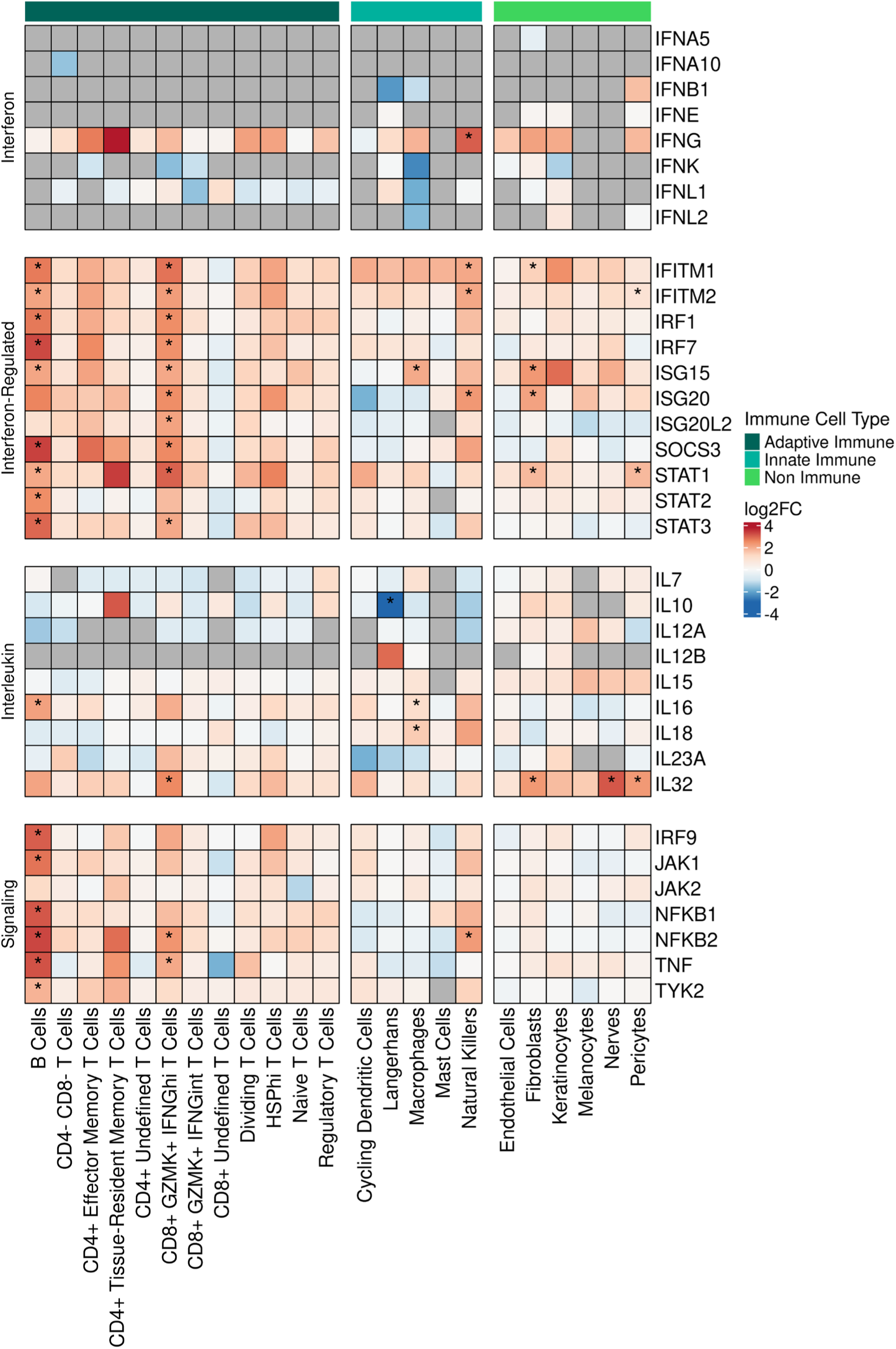
Gene expression of interferon, interferon-regulated, interleukin, and signaling-related genes in the skin. Heatmap showing the average fold change of the expression of select genes across the full set of skin cell types. The bar on the top indicates whether or not the cell types plotted are immune-related (dark green for adaptive immune cell types, cyan for innate immune cell types, and bright green for non-immune cell types). Within the heatmap itself, red indicates upregulation, and blue indicates downregulation, with asterisks denoting statistically significant changes (FDR < 0.05). Gray cells indicate no expression of the relevant gene for that cell type (not applicable). Differential expression was calculated between the EM and control portions of each cell type.

To determine which interferons might be produced locally in the skin and subsequently drive the IFN response signature, we carried out a differential gene expression analysis of EM versus control skin in every cell subset (Figure 7, Supplemental Figure 9). Out of all of the interferon genes, only IFNG was significantly upregulated (and only in NK cells); most other cell types exhibited increased IFNG expression but at levels that did not achieve statistical significance. The only type I IFNs that were detected as differentially upregulated (albeit not significantly) were IFNB1 in pericytes and IFNK in fibroblasts.

### Expression of select cytokines is upregulated in EM lesions

Finally, we investigated the expression of interleukin genes to identify which cells in the EM contribute to cell recruitment through cytokine production (Figure 7, Supplemental Figure 10). Macrophages significantly upregulated IL16 and IL18, which are known to induce IFNG expression and are consistent with an M1 phenotype (54). IL16, which is also a chemoattractant for CD4^+^ T cells, was significantly upregulated in B cells as well. None of the T cell subsets showed increased expression of any interleukin genes, except for CD8^+^ GZMK^+^ IFNG^hi^ T cells which upregulated IL32, a cytokine that can promote FOXP3^+^ Treg cell development and influence dendritic cell function to promote proinflammatory T cell responses (55, 56). IL32 was also significantly upregulated within the fibroblasts, nerve cells, and pericytes. These findings suggest that a few select cell types expressing key interleukins drive interferon production and modulate the recruitment and function of immune cells.

## Discussion

The skin represents the first barrier to infection by arthropod-transmitted pathogens and is the site where *Bb* spirochetes must establish infection and persist to complete their life cycles (57). Transmission of *Bb* to the mammalian host is preceded by replication, programmatic changes in gene expression, and migration from the *Ixodes* tick midgut to the salivary glands prior to deposition into the dermal layer of the skin at the bite site. Tick salivary protein interactions with newly expressed *Bb* proteins and the pharmacologic properties of tick saliva inhibit local immunity to permit *Bb* to successfully establish infection in the dermis (58). These actions of tick saliva as well as the introduction of spirochetes into the skin are likely to perturb the bite site microbiome (58). The EM lesion temporally appears after tick detachment when the local effects of tick saliva are expected to be waning, and *Bb* (which is an extracellular bacterial pathogen) begins to migrate outward through the collagen-rich dermal connective tissue. By the time EM is brought to medical attention days to weeks after a tick bite, histopathology of the lesion reveals the dominance of T cells, macrophages, and DCs as the principal immune cells responding to the introduction of *Bb* into the complex microbial community of the bite site (32).

Earlier studies have characterized the immune cell infiltrates in EM lesions by *in situ* hybridization techniques (14), flow cytometry of cells along with cytokine analysis of fluid extracted from EM lesions (15), and more recently through transcriptomic analysis of EM skin biopsies (9, 19). These techniques have noted the aforementioned predominance of T cells and macrophages along with the presence of elevated levels of inflammatory (IFNG, IL1B, IL6, TNFA) and anti-inflammatory (IL10) cytokines in EM lesions. Phenotypic analysis of the T cell subsets from fluid in blisters experimentally induced over the EM lesion revealed that the proportion of CD4^+^ T cells (68%) was much higher than CD8^+^ T cells (20%), with the CD4^+^ T cell population dominated by CD27^-^ CD45RO^+^ memory effector cells and strongly polarized toward a Th1 phenotype (15). A transcriptomic analysis via microarray on whole skin digests of the EM lesion supported some of these findings and also revealed a potential role for the kynurenine pathway and tryptophan metabolites in the regulation of the inflammatory response (9). A model for the cutaneous immune response based on transcriptomic microarray analysis has been proposed supporting the viewpoint that innate immune cells (macrophages and DCs) responding to *Bb* inflammatory components recruit CD4^+^ Th1 cells that result in the strong IFNG signatures seen in the EM lesion (9).

Our studies using scRNA-Seq combined with TCR sequencing on whole EM skin cell digests have provided new information that refine our understanding of the host response to *Bb* in the skin. Our results reveal enrichment of many immune cell subsets in the EM lesion along with the expected dominance of T cells. CD8^+^ T cells constituted the largest T cell population, with both CD8^+^ GZMK^+^ IFNG^+^ T cell subsets enriched in the EM lesion and contributing to the IFNG response, as well as possibly including TEM_RA_ cells. Although we do not have corresponding protein expression to corroborate expression of CD45RA (which is known to be re-expressed on TEM_RA_ cells), our analysis of CD45 gene splicing detected CD45RA mainly within the CD8^+^ GZMK^+^ IFNG^int^ T cell subset rather than the CD8^+^ GZMK^+^ IFNG^hi^ subset, indicating that cells that could be classified as TEM_RA_ are more prevalent in this subset. TEM_RA_ cells can have effector functions such as cytotoxicity and are associated with cell senescence in aging (59, 60); however, we observed an increased proportion of CD8^+^ GZMK^+^ IFNG^+^ T cells in EM lesions from all study participants (age range 31-76, median 68), indicating that their presence was not a function of aging per se.

Furthermore, we found that not all CD8^+^ GZMK^+^ IFNG^+^ cells expressed characteristic cytotoxic genes. A greater proportion of the CD8^+^ GZMK^+^ IFNG^hi^ T cells contained cells with low or undetectable levels of GZMB and PRF1, in comparison to the CD8^+^ GZMK^+^ IFNG^int^ T cells. These CD8^+^ GZMB^low^ PRF1^neg^ T cells are reminiscent of CD8^+^ GZMK^+^ T cells (T_teK_ cells) found in rheumatoid arthritis synovium and at barrier sites, where they have been proposed to initiate or sustain inflammatory responses (33, 61). CD8^+^ GZMK^+^ T cells can be activated in an antigen-independent fashion in response to cytokines alone and in this way share features of innate immune cells (33, 62). Although they are enriched in peripheral non-lymphoid tissues, they can also be detected in the circulation (63). We found that both CD8^+^ GZMK^+^ IFNG^hi^ T cells and CD8^+^ GZMK^+^ IFNG^int^ T cells exhibited clonal expansion in the EM lesion and subsets within both populations expressed TCRs that were clonally related to T cells in the blood. Notably, some clonotypes were also shared between the two CD8^+^ GZMK^+^ IFNG^+^ T cell populations in the EM lesions. CD8^+^ GZMK^+^ IFNG^hi^ T cells in the control samples, however, do not share any clonotypes with blood T cells, in contrast to CD8^+^ GZMK^+^ IFNG^int^ T cells, in which a small number of clonotypes in the control skin were shared with the blood and/or the EM lesions. Taken together, these results suggest that the two populations of CD8^+^ GZMK^+^ IFNG^+^ T cells are related to one another in both the EM lesions and the blood, but that only CD8^+^ GZMK^+^ IFNG^int^ cells from the blood and EM lesions also have clonal relatives in the control skin. However, a comparison of the genes differentially expressed by CD8^+^ GZMK^+^ IFNG^hi^ T cells versus the CD8^+^ GZMK^+^ IFNG^int^ T cells revealed substantial differences, with the former including genes associated with inflammation and its regulation and the latter including genes associated with cytotoxicity. The kinetics of the responses of these subsets are not known, but together they may represent cells that respond to disruption of the skin barrier and introduction of *Bb* while maintaining a cytotoxic population that keeps the cutaneous microbiome in check. Additional research beyond the scope of this study would be needed to further define the relationships of the different CD8^+^ GZMK^+^ IFNG^+^ T cell populations between the two tissue sites and the blood.

Our gene set enrichment analysis identified both Type I and Type II IFN pathways, along with other immune pathways involved in host defense and immune regulation as significantly upregulated in the EM lesion. The finding that B cells and CD8^+^ GZMK^+^ IFNG^hi^ T cells exhibited robust differential expression of IFN-stimulated genes suggests that both cell types may play important roles in the early host defense and contribute to the *Bb*-specific CD4^+^ T cell responses. B cells in particular upregulated the T cell recruiting cytokine IL16 that promotes CD4^+^ Th1 cell differentiation whereas CD8^+^ GZMK^+^ IFNG^hi^ T cells upregulated expression of IL32, a cytokine important for immune responses against viruses and intracellular bacteria (54–56). In assessing chemokine expression, we noted that fibroblasts, not macrophages, differentially expressed the broadest array of T cell recruiting chemokines. Fibroblasts are the major component of the dermis and comprise diverse subtypes, including those involved in homeostasis of the extracellular matrix and others that have a role in immune responses, inflammation, and host defense against pathogens (reviewed in (63)). *In vitro* studies using microarrays to assess gene expression have documented that primary human dermal fibroblasts exposed to *Bb sensu stricto* spirochetes upregulate expression of several cytokines and chemokines, including IL6, IL8, CCL2, CCL5, CXCL1, CXCL2, CXCL6, CXCL9, and CXCL10 (64). These results have led to the hypothesis that dermal fibroblasts responding to the presence of *Bb* in the skin play a key role in recruiting immune cells to the infection site. Our study provides support for this hypothesis by showing the significant increase in gene expression of CCL4, CCL5, CCR1, CCR5, CXCL9, CXC10, and CXC11 by fibroblasts in EM lesions. In addition, we found upregulation of chemokine genes by other non-immune cells such as endothelial cells and pericytes, which may serve to recruit T cells and macrophages to perivascular tissues consistent with the histopathology characteristic of EM lesions.

While we have evidence that fibroblasts recruit inflammatory cells to the EM lesion, they also may contribute to regulation of the inflammatory response. We found that fibroblasts were the only skin cell type that exhibited significant differential expression of IDO1 (Supplemental Figure 9), the gene for a rate-limiting enzyme for tryptophan catabolism to kynurenine (65) that was previously found to be upregulated in transcriptomic microarray analyses of EM digests (9). Tryptophan depletion can suppress T cell proliferation, and genes in the kynurenine pathway can induce apoptosis of Tregs and other T cells through activation of the aryl hydrocarbon receptor (66). KMO and KYNU, two other genes associated with tryptophan catabolism, were differentially expressed within the B cell population, suggesting that the kynurenine pathway was activated, which could potentially lead to apoptosis of B cells (67) thereby interfering with the B cell defense in the skin.

While the immune changes we observe in the EM lesion may be triggered by *Bb* infection, they may also reflect in part ongoing immune responses that arise as a result of tick feeding and disruption of the skin microbiome. European studies that examined the cutaneous immune response to naturally acquired bites from *Ixodes ricinus* ticks that did not harbor *Borrelia spp.* spirochetes have documented an increasing presence of lymphocytes over time (13, 68). Flow cytometry analyses of these lymphocytes identified increases in both B and T cells, with a decrease in the CD4^+^ to CD8^+^ T cell ratio in comparison to uninvolved skin, and an increase in the tissue-resident memory cells. A gene expression analysis performed on skin biopsies from people bitten by *Borrelia spp.-*infected vs *Borrelia* spp.-uninfected *Ixodes ricinus* ticks has recently been reported (69). Skin samples were taken from the tick bite site as well as uninvolved skin 7-10 days after the tick was removed. Although the study did not find statistically significant differences in gene expression between the two groups, Ingenuity Pathway Analysis performed on data from the *Borrelia spp.-*infected tick group vs autologous skin revealed elevation in Type I IFN pathways and regulators of T cell and NK cell responses. Taken together, these findings demonstrate that the act of tick feeding elicits changes in the immune cell repertoire of the skin, with an increase in B cells and T cells, the latter of which reveal skewing toward increased CD8^+^ T cell responses. While our skin samples were taken at the leading edge of the EM lesion, away from the bite site, it is possible that some of the changes we observe in the immune cell populations result from tick feeding and alterations of the microbiome (68).

There is limited information on the role of CD8^+^ T cells in LD in humans, and no prior study has phenotypically or functionally characterized this population in the EM lesion. A European study examining TCRVβ chain usage of T cells isolated from cerebrospinal fluid (CSF) and peripheral blood of people presenting with neuroborreliosis identified clonal expansion of CD8^+^ T cells within the CSF that had a memory phenotype (CD45RO^+^, CD28^+^) and that decreased in number after treatment (70). These cells expressed CD69 and CCR5, produced IFNG after *in vitro* stimulation with CD3/CD28 beads, and released TNFA and to a lesser extent IFNG when incubated with DCs and *Bb* lysate. However, they did not exhibit cytotoxicity when stimulated with EBV-transformed B cells pulsed with *Bb* lysate. One proposed interpretation of these findings was that the CSF CD8^+^ T cell clonotypes were helping to initiate the inflammatory response to *Bb* infection in the CNS without exerting cytotoxic effects on antigen-presenting cells that may be activating them. Other studies have identified expansion of cytotoxic CD8^+^ T cells in blood samples from people experiencing protracted symptoms after neuroborreliosis when their PBMCs were cocultured with macrophages pre-exposed to live spirochetes (71). *Bb* antigen-specific cytotoxicity has also been observed in CD8^+^ T cell lines derived from peripheral blood and synovial fluid of people with Lyme arthritis, and notably T cells exhibiting this phenotype were only detected in peripheral blood after resolution of arthritis (72). A more recent study found a greater percentage of CD8^+^ T cells in the peripheral blood of people who had post-infectious Lyme arthritis in comparison to those who resolved joint inflammation with antibiotic therapy (73). In contrast to rheumatoid arthritis, in which the CD8^+^ T cells had low levels of granzyme B and perforin (33), CD8^+^ T cells found in Lyme arthritis samples were characterized by elevated levels of these cytotoxic molecules. Given the prevalence of CD8^+^ T cells near vasculature on immunohistochemistry, it was postulated that cytotoxic CD8^+^ T cells may lead to microvascular changes that are characteristic of post-infectious Lyme arthritis. Taken together with our results identifying CD8^+^ GZMK^+^ IFNG^+^ T cell subsets that are predicted to have low cytotoxicity based on transcriptomic gene expression, CD8^+^ T cells in LD may comprise distinct subsets, one capable of promoting an inflammatory state early in *Bb* infection in the skin or CNS while minimizing tissue damage, and others with cytotoxic potential contributing to adverse sequelae.

In summary, our single cell analysis has identified CD8^+^ GZMK^+^ IFNG^+^ T cells enriched in the EM lesion that may play an important role in the immune response at the skin barrier site to Ixodes tick-transmitted *Bb* infection. Conclusions from our study may be limited by the relatively small number of participants with skewing towards older individuals and women. In addition, we postulate that CD8^+^ GZMK^+^ IFNG^+^ T cells, along with B cells, instructed by nonimmune cells such as fibroblasts, shape the cellular response to the introduction of *Bb* and the concomitant perturbation of the cutaneous microbiome. Although we have focused on the tick-transmitted bacterium *Bb*, our findings may have relevance to the cutaneous host defense against other vector-borne pathogens. Additional studies to assess specific interactions among immune and nonimmune cells, such as spatial transcriptomics together with protein expression analyses, could further enhance our understanding of the initial host defense against vector-borne pathogens.

## Methods

### Sex as a biological variable

Male (n = 4) and female (n = 9) participants were recruited, and similar findings were reported for both sexes. Sex and age were not considered as analysis variables.

### Subject recruitment and sampling

Thirteen participants ranging from 31 to 76 years of age were recruited from study sites affiliated with Yale School of Medicine/Yale New Haven Hospital in New Haven, CT, and from Mansfield Family Practice in Storrs, CT. Eleven participants met the 2017 CDC criteria for Lyme disease (74) and had at least one EM lesion. Two additional subjects with 4 cm EM lesions had probable LD based on physician-diagnosed EM in the setting of known tick bites, one of whom had positive Lyme serologies. Participants were enrolled during the period of May to October in 2019 (n = 6), and in 2020 (n = 7). Clinical details of the participants recruited in 2019 have been previously reported (see dataset 1 in Jiang et al. (19)), and all participants enrolled in this study are described in Table 1.

### Sample processing

Blood and skin samples were collected from all participants and processed for scRNA-Seq and BCR and TCR sequencing as previously described (19). Whole blood samples were collected in separate blood collection tubes for sera, DNA extraction, and PBMC isolation. Ficoll-Paque PLUS purified PBMCs isolated as described previously (19) were cryopreserved in 90% human AB negative sera (GeminiBio) and 10% DMSO for bulk BCR and TCR sequencing. Three mm skin punch biopsies were obtained from the leading edge of the EM lesion and from uninvolved (control) skin at least 2 cm away from the EM lesion. Biopsies were placed into MACS Tissue Storage Solution (Miltenyi Biotec, 130-100-008) for transport to the laboratory where they were immediately processed into single cell suspensions using the Whole Skin Dissociation Kit (Miltenyi Biotec, 130-101-540) according to the manufacturer’s recommendation, as previously described (19). Library construction from single cell suspensions was performed using the 10x Genomics Chromium Single Cell 5’ Reagent Kit (both gene expression and immune profiling) and sequencing was done on the Illumina MiSeq platform. Sample processing produced three datasets; one representing samples collected in 2019 and the other two representing samples collected in 2020.

### scRNA-Seq data processing and quality control

The raw sequencing data from all of the datasets were processed with 10x Genomics’ Cell Ranger v6.1.2 using GRCh38-2020-A as the GEX reference and vdj_GRCh38_alts_ensembl-5.0.0 as the VDJ reference. To have greater control over quality control decisions, *cellranger count* and *vdj* were run separately instead of *cellranger multi*. Upon checking the average read depths across the blood and skin datasets, the gene expression data was rerun using *cellranger aggr* to account for significant differences among datasets (dataset 1 had an average of 132 million reads, dataset 2 had an average of 198 million reads, and dataset 3 had an average of 390 million reads) (Supplemental Figure 11). This subsampled every GEX sample down to the lowest recorded read depth. An average of 5,882 cells were identified per sample and there were an estimated 41,568 cells in the blood and 146,662 cells in the (combined EM and control) skin (20).

### Gene expression analysis

The count matrices were processed using Seurat v4.4 as the single cell ecosystem. After filtering out low-quality cells by removing cells with fewer than 200 genes and mitochondrial content greater than 15%, there were 40,733 cells remaining in the blood and 136,002 cells remaining in the skin (87,836 from the EMs and 48,166 from the controls). The data were then log-normalized with a scale.factor of 10,000 using *NormalizeData.* The top 2000 most highly variable genes were identified using the *vst* method of *FindVariableFeatures* and linearly scaled with *ScaleData.* IG (immunoglobulin) and TR (T cell receptor) genes were removed from the variable features using Ensembl 93 as a reference for the gene names. Principal component analysis was run on 30 PCs with *RunPCA* and then graph-based clustering was conducted on 20 dimensions (chosen based on the inflection point of an elbow plot) using *FindNeighbors* for shared nearest neighbors and *FindClusters* for Louvain-based community detection. Non-linear dimensionality reduction for visualization was done using *RunUMAP*. Default parameters were used for all functions unless otherwise specified. The blood and skin datasets both showed adequate mixings of samples, indicating no major batch effects and therefore no need to add a computational integration (harmonization) step.

Clusters were computed and annotated using known canonical marker genes (Supplemental Figures 1 and 2). The clustering resolution was determined through iteration and refinement, with a final resolution of 1.0 being selected for the skin and 0.4 for the blood (Figures 1, 2, and Supplemental Figure 12). We chose to initially over-cluster with the understanding that similar clusters could then be combined with the same cell type label. Some clusters may contain mixtures of cell types. Cell type annotation was further supported by integrating TCR information with plots such as DotPlots to better identify T cells in both tissue types (Supplemental Figures 1 and 2). As 10x Genomics does not currently support the detection of gamma-delta chains, we used the combination of several other approaches to define a gamma-delta T cell cluster within the subclustered blood T cell data; reconstruction of TCRs from GEX data using TRUST4 (75), automated annotation using Azimuth (76) and expression of specific marker genes (TRDV2, TRGV9, TRGV10).

Doublets were calculated on the level of Seurat clusters per sample and removed using scDblFinder (77). After testing several rates, 0.01 was chosen as the expected doublet rate following 10x Genomics documentation (1% per 1000 captured cells). There were 38,942 cells remaining in the blood and 123,304 cells remaining in the skin (83,860 from the EMs and 39,444 from the controls) post-doublet removal. The majority of doublets removed from the blood came from cells labeled as DCs and T cells and from the skin came from cells labeled as fibroblasts and keratinocytes.

We identified possible TEM_RA_s using IDEIS (identification of isoforms), a recently developed tool for assessing differentially spliced variants of CD45 in 10x Genomics data (78). It works by creating a transcriptome reference from the raw BAM files and mapping reads to the relevant exons. 17% of the skin T cells had reads successfully mapped, which is within the range predicted by the IDEIS authors based on reads per cell.

### Immune profiling analysis

Paired-end FASTQ reads of single cell TCR data were processed using the *cellranger vdj* command from 10x Genomics Cell Ranger v6.1.2 (20) for alignment against the human reference. Sequences were further processed using the nextflow pipeline nf-core/airrflow v4.1.0 (79), which uses tools from the Immcantation pipeline (80, 81). airrflow performed V(D)J germline gene reassignment using IgBLAST v1.20.0 and the IMGT database (82, 83). Only productively rearranged sequences with valid V and J gene annotations and junction lengths divisible by 3 were retained. Cells with only alpha chains and no paired beta chains were discarded.

Bulk TCR data were processed as previously described (19) and combined with single cell TCR data to perform clonal lineage analysis. TCR clonotypes from each participant were defined as sequences sharing the same V and J genes and an identical junction region in the beta chains, using the *identicalClones* function from the SCOPer package (v1.2.1) (84). Clonal overlaps were visualized using ggVennDiagram v1.5.2 (85) and clonal expansion was visualized using APackOfTheClones v1.2.4 (86).

### Subclustering

Subclustered T cell populations were defined for the blood and the skin in several stages. In the first stage, cells already annotated as T cells on the full tissue level (Figure 1) were selected along with cells with paired TCRs, as the latter may have been real T cells that simply did not have clear enough expression to be annotated as such. After redoing the gene expression processing steps above (including removing the IG and TR genes from variable features again and reclustering), clusters that did not appear to be T cells based on expression of canonical T cell marker genes were removed. In the second stage, the now uncontaminated T cells were reclustered (with a resolution of 1.5 for both the blood T cells and the skin T cells) and annotated with T cell subtypes using marker genes again.

For analysis of the full skin (e.g. for differential gene expression analysis), cells from the original “T Cells” group were renamed with the identified T cell subtypes when possible. Any cells from the “T Cells” group that were not selected for subclustering were removed as we did not believe that they were truly T cells.

### Differential gene expression analysis and gene set enrichment analysis

To identify differentially expressed genes between control and EM samples across skin cell types, we conducted a pseudobulk analysis by aggregating cell counts for each cell type within each sample and applying DESeq2 (87). Genes were considered to be significantly differentially expressed using an FDR threshold of 0.05. Heatmaps were created using ComplexHeatmap v2.22.0. Gene set enrichment analysis (GSEA) was performed on the ranked gene list from the pseudobulk analysis on the Hallmark gene sets from the Molecular Signature Database (MSigDB) (v2024.1.Hs) (88) using ClusterProfiler v4.14.4 (89). We specifically looked for gene sets that were enriched among the upregulated genes in EM (i.e. positive enrichment scores), with significance determined by a Bonferroni-corrected p-value less than 0.01.

### Statistical analysis

All analyses were done with R v4.3. The significance threshold was 0.05 for FDR-adjusted p-values unless otherwise stated. A paired Wilcoxon signed-rank test was used for comparing the subset frequencies between control and EM across subjects. A two-sample test for equality of proportions with continuity correction was used to compare clonal overlaps between the blood and the EM with the blood and the control skin.

### Study approval

This research was conducted under human research protocols approved by the IRB of Yale University. Written informed consent was received from all participants prior to inclusion in the study.

## Data availability

The code used for analysis is made available at https://github.com/Kleinstein-Lab/LymeTCells. The raw and processed scRNA-Seq and scTCR-Seq data used in this publication have been deposited in NCBI’s Gene Expression Omnibus (GEO) (90) and are accessible through the GEO Series accession numbers GSE297325 (https://www.ncbi.nlm.nih.gov/geo/query/acc.cgi?acc=GSE297325) and GSE169440 (https://www.ncbi.nlm.nih.gov/geo/query/acc.cgi?acc=GSE169440).

## Author contributions

LKB, AAB, and SHK designed the research studies. LKB and AAB oversaw participant recruitment and tissue sample collection. All authors participated in data acquisition and analysis, with EA, HM, and PF performing bioinformatics analysis. EA, LKB, HM, AAB, and SHK wrote the manuscript, with all authors editing and approving the final content for submission.

## Acknowledgments

This project was supported by the National Institute of Allergy and Infectious Diseases of the NIH through the U19-AI089992 and T15LM007056 grants. It was also supported by the Harold W. Jockers Professorship to LKB.

We thank Kenneth R. Dardick for help with participant recruitment and all of our participants for contributing to this study and for donating tissue samples. We thank Jialing Mao, Ming Li, and the Yale Center for Genome Analysis for processing the samples, as well as Ruoyi Jiang and Khadir Raddassi for technical assistance. We also thank Mary Tomayko for her expertise in skin cell types, Donna Farber, Peter Sims, and Bjoern Peters for their insights into T cell subtypes, and Gur Yaari for advice on bioinformatics analysis. We thank Edward Lee for his help with running TRUST4 and Wengyao Jiang for her assistance in submitting our data to the GEO repository and making the data accessible to the research community.

## Abbreviations and acronyms

Bb: Borrelia burgdorferi
BCR: B Cell Receptor
DC: Dendritic Cell
EM: Erythema Migrans
FDR: false discovery rate
GEO: Gene Expression Omnibus
HSP: Heat Shock Protein
IFN: interferon
IRG: Interferon-Regulated Gene
LD: Lyme Disease
NK: natural killer [cells]
PBMC: peripheral blood mononuclear cell
TCR: T Cell Receptor
TEMRA: T effector memory cells re-expressing CD45RA
Tr1: Type 1 regulatory T cells

## Supplemental Figures

**Supplemental Figure 1.**
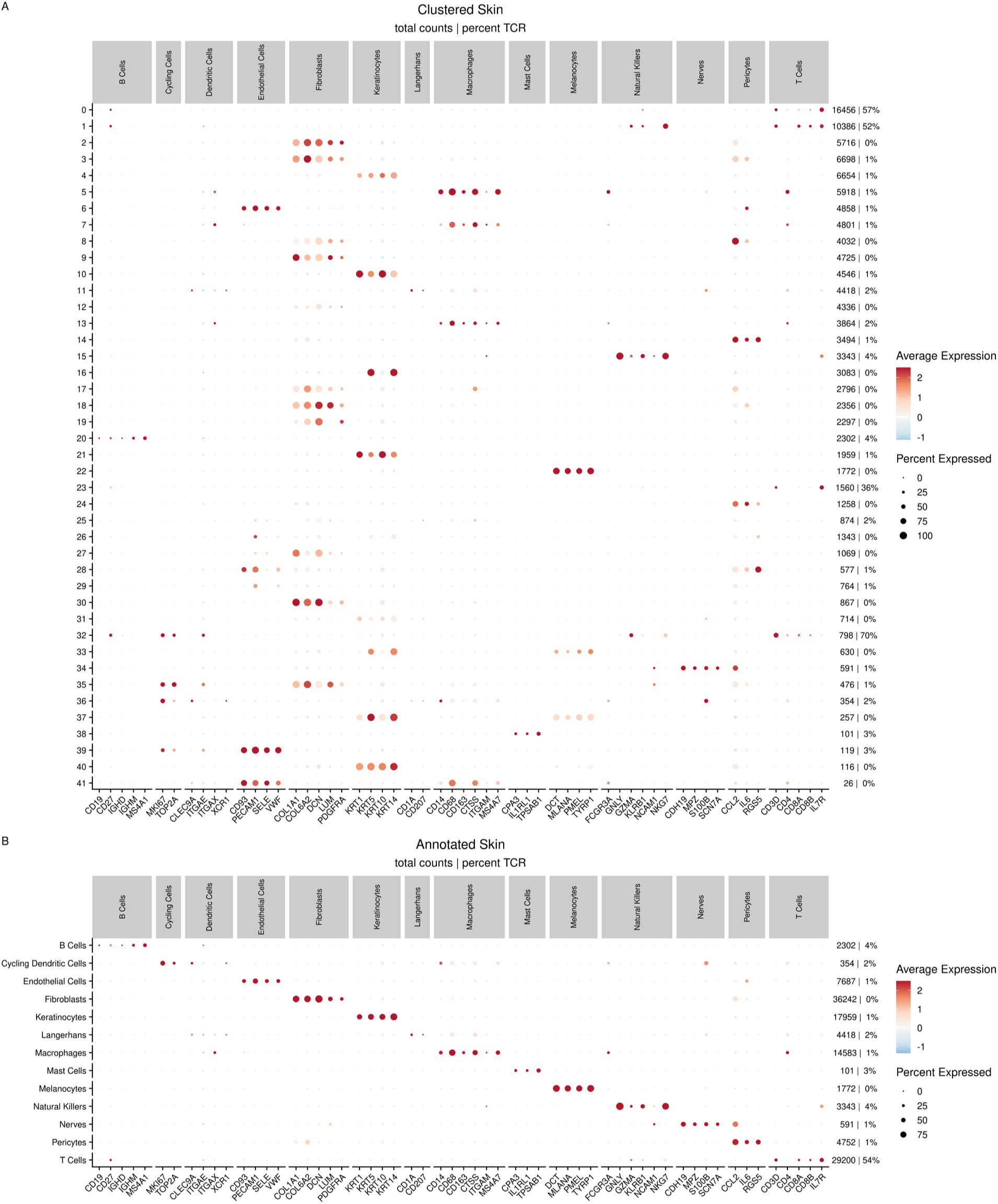
Clustering and annotation of the skin. **A)** Dot plot showing the average expression of characteristic marker genes for each cluster within the skin. **B)** Dot plot showing the average expression of the same set of markers for each annotated cell type within the skin. Red indicates higher levels of expression and blue indicates lower levels of expression, with the size of the dot reflecting the percentage of that cluster of cell type that is expressing the corresponding marker. Numbers in the two columns on the right side of the plots indicate the total number of cells in that cluster or cell type and the percentage that have a paired TCR, respectively.

**Supplemental Figure 2.**
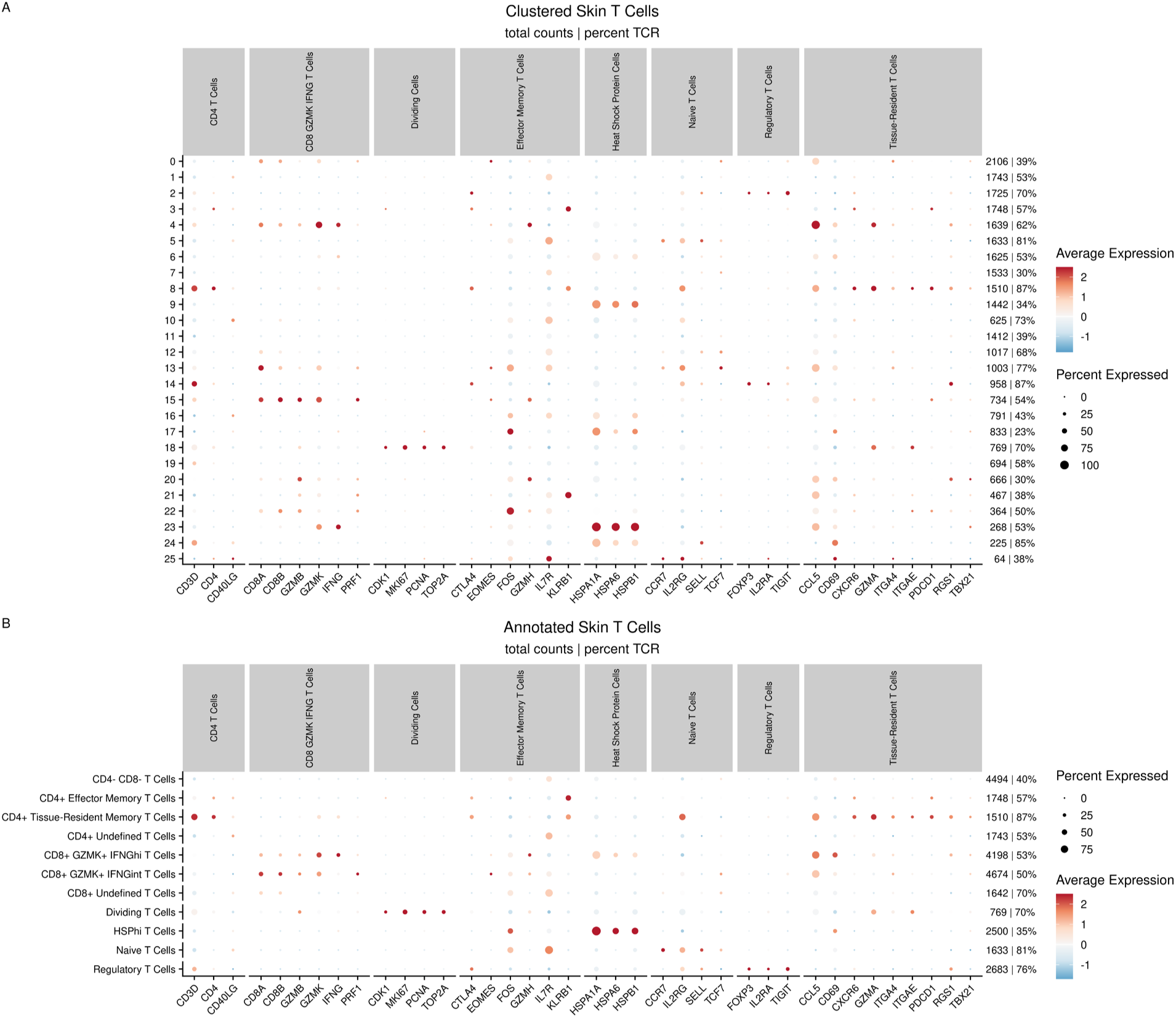
Clustering and annotation of the skin T cells. **A)** Dot plot showing the average expression of characteristic marker genes for each cluster within the skin T cells. **B)** Dot plot showing the average expression of the same set of markers for each annotated cell type within the skin. Red indicates higher levels of expression and blue indicates lower levels of expression, with the size of the dot reflecting the percentage of that cluster of cell type which is expressing the corresponding marker. The two columns of information on the right side of the plots indicate the total number of cells in that cluster or cell type and the percentage which have a paired TCR respectively.

**Supplemental Figure 3.**
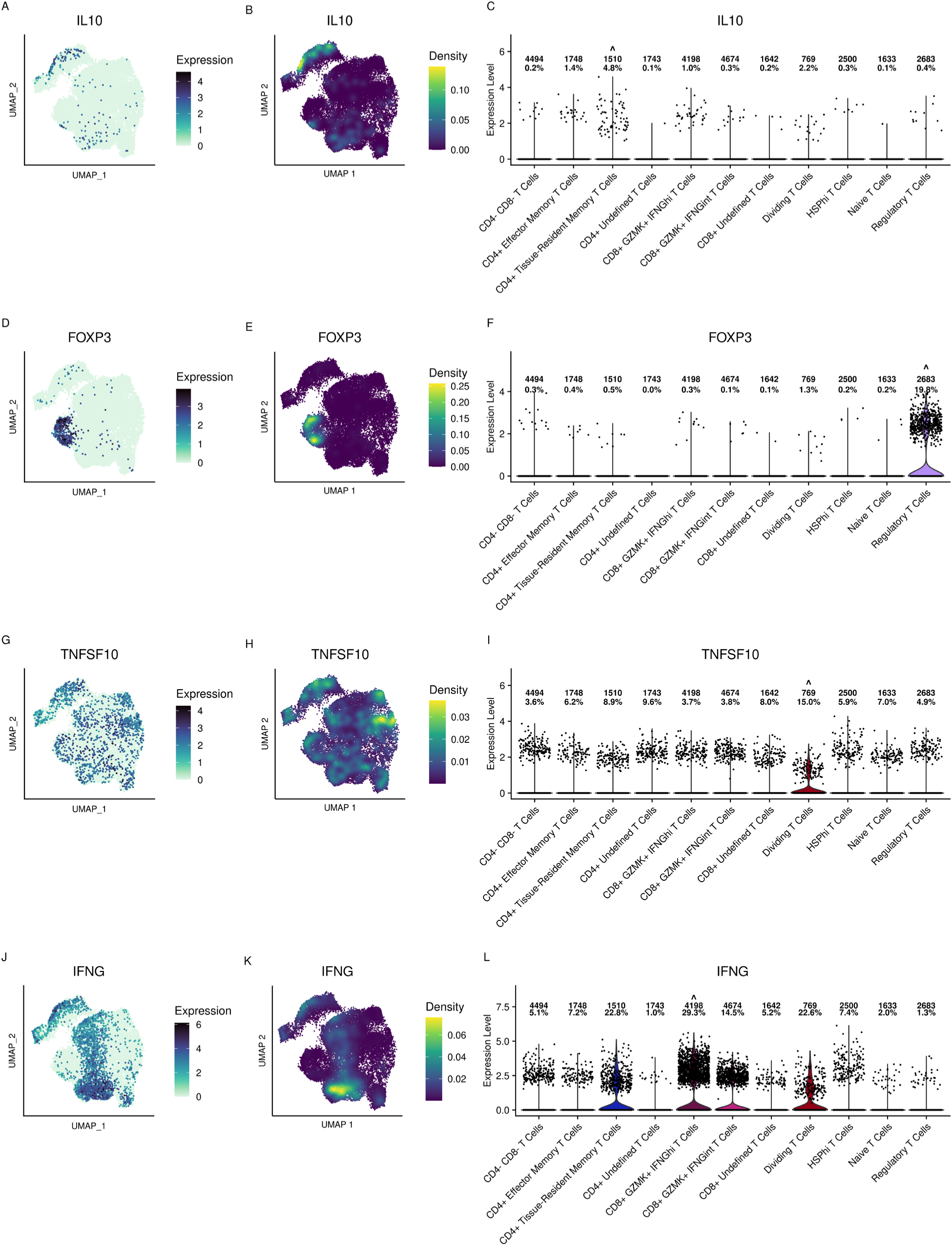
Expression of select genes within the skin T cells. Each row shows the expression of IL10 (**A-C**), FOXP3 (**D-F**), TNFSF10/TRAIL (**G-I**), and IFNG (**J-L**) respectively. **A)**, **D)**, **G)**, and **J)** show the average expression of each gene upon the skin T cells UMAP, with darker colors corresponding to higher expression. **B)**, **E)**, **H)**, and **L)** show expression as a feature of kernel density using the Nebulosa package to allow for better visualization. **C)**, **F)**, **H)**, and **L)** show expression on a cell type level, with points representing individual cells. The top row of numbers within the violin plots is a count of how many cells are in that cell type, and the second row is the percentage of them with positive expression of the relative gene.

**Supplemental Figure 4.**
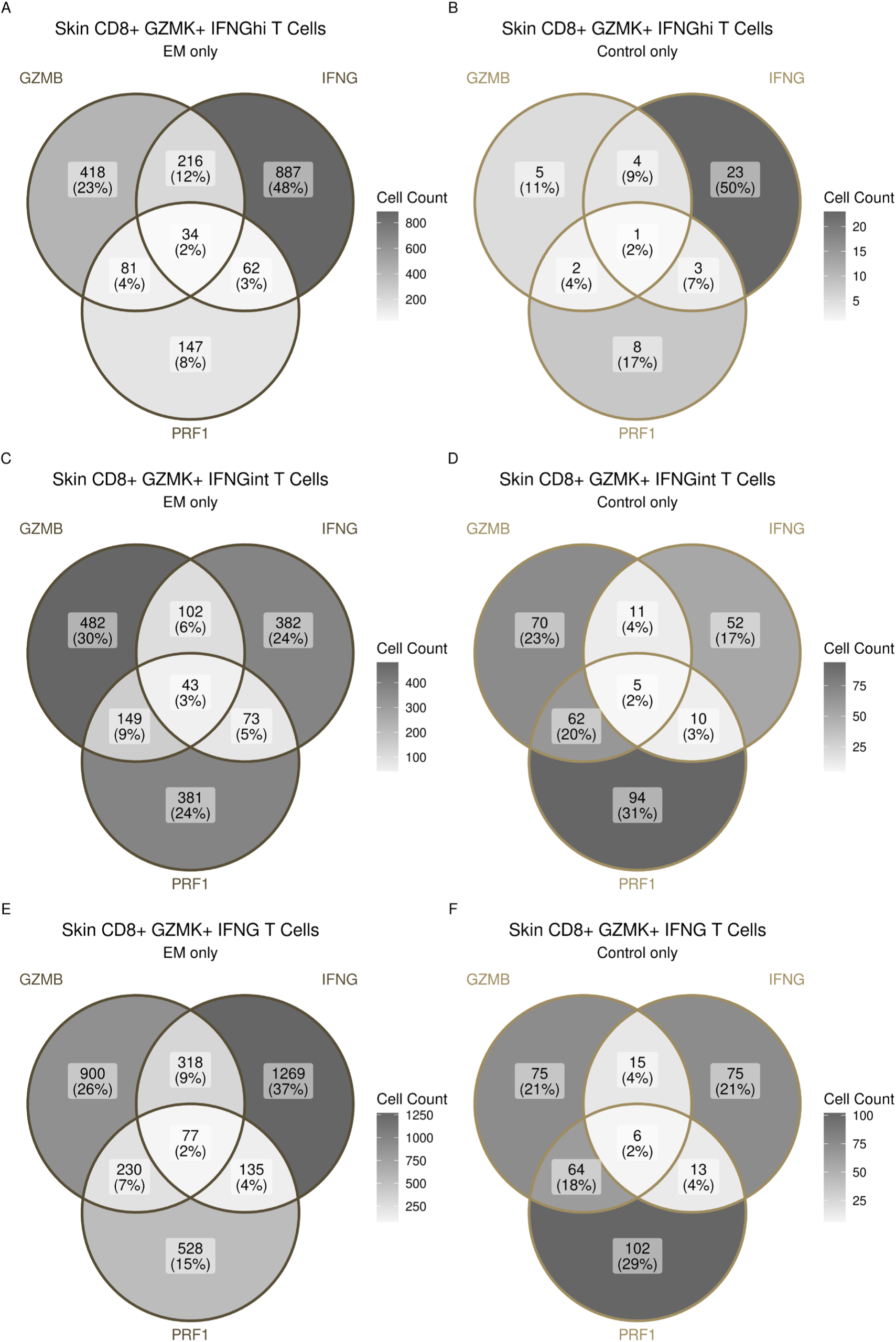
Positive expression overlaps of GZMB, IFNG, and PRF1 within the CD8+ GZMK+ IFNG+ T cells. Each Venn diagram represents the number of cells with positive expression of GZMB, IFNG, and/or PRF1, with percentages relative to the total cells being represented in that Venn diagram. The left column is for only the EM subset and the right column is for only the control subset. **A)** and **B)** are for the CD8+ GZMK+ IFNGhi T cells, **C)** and **D)** are for the CD8+ GZMK+ IFNGint T cells, and **E)** and **F)** are for both cell types combined.

**Supplemental Figure 5.**
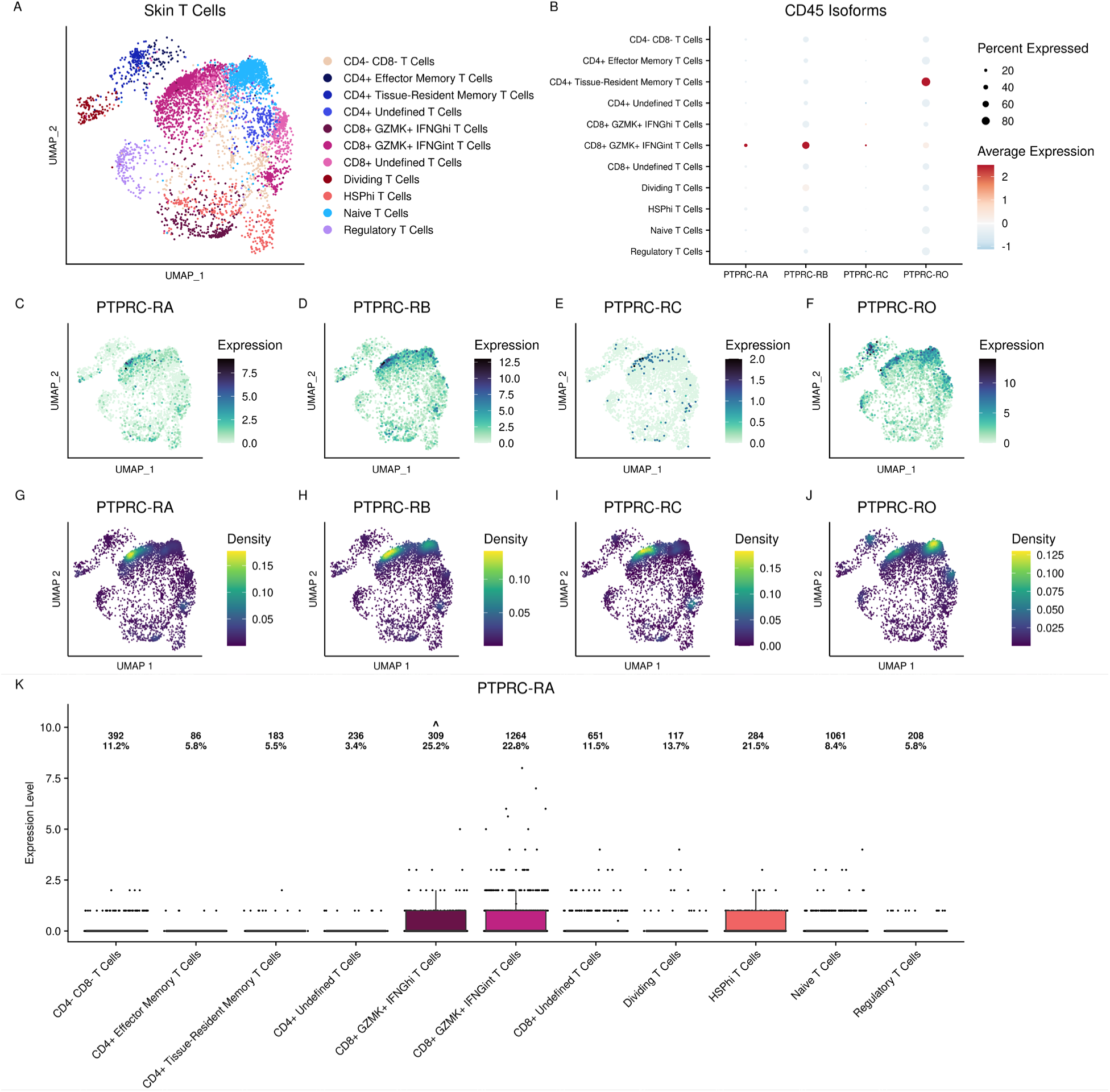
CD45 spliced isoforms in skin T cells using IDEIS. **A)** UMAP showing all of the skin T cells that had a spliced isoform of CD45 (RA, RB, RC, or RO) detected with the IDEIS package. **B)** Dot plot showing the expression of each isoform per T cell subtype. **C-F)** Expression of each isoform upon the skin T cells UMAP, with darker colors corresponding to higher expression. **G-L)** Expression as a feature of kernel density using the Nebulosa package to allow for better visualization. **K)** Expression on a cell type level, with points representing individual cells. The top row of numbers within the box plot is a count of how many cells are in that cell type, and the second row is the percentage of them with positive expression of CD45RA.

**Supplemental Figure 6.**
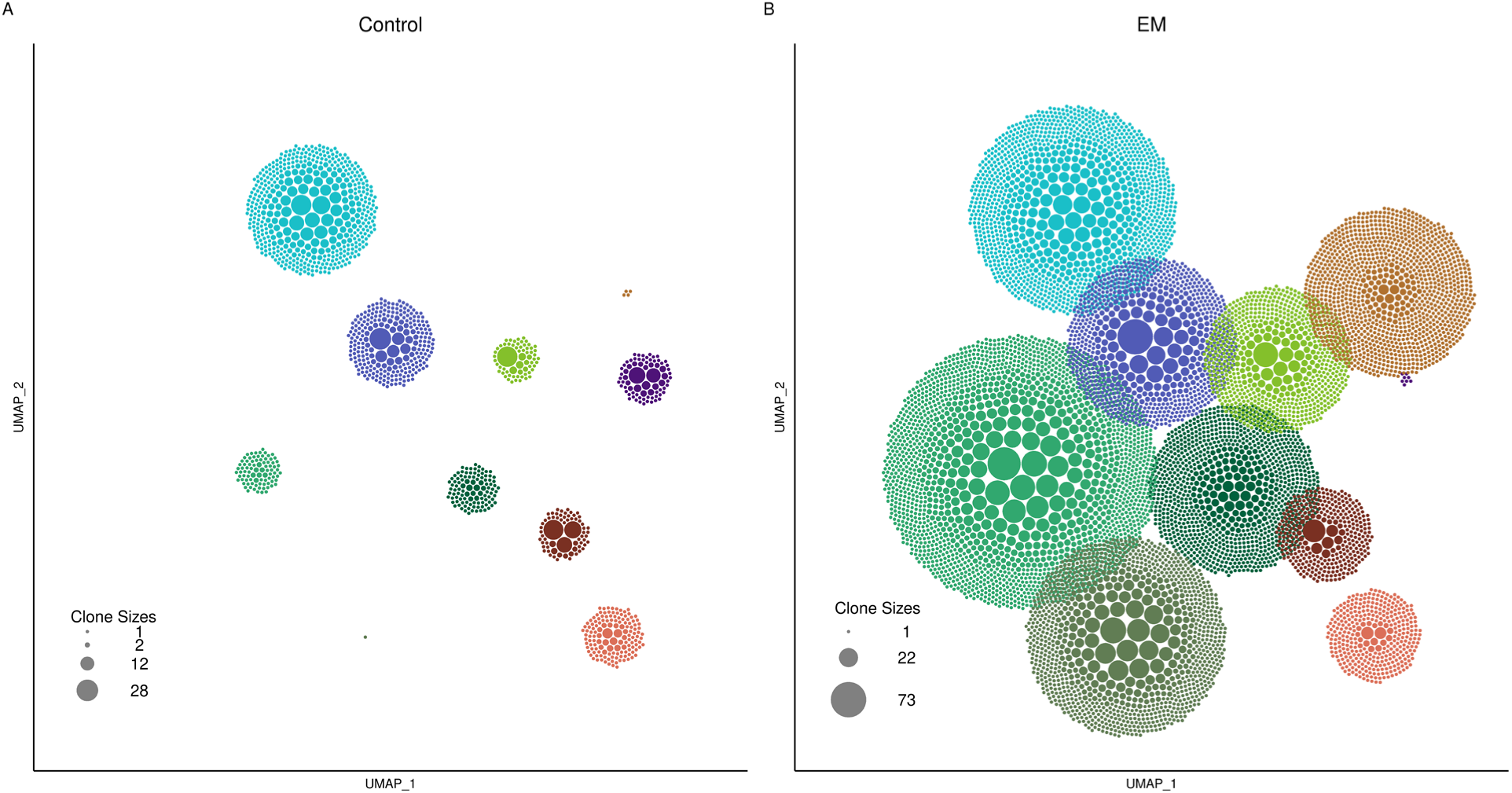
Clonal expansion of skin T cells. **A)** and **B)** UMAP of the skin T cells in control and EM, respectively, with points representing clonotype sizes within each participant. The scale of the point size is consistent for both sample types. Participants without a paired control and EM are not shown.

**Supplemental Figure 7.**
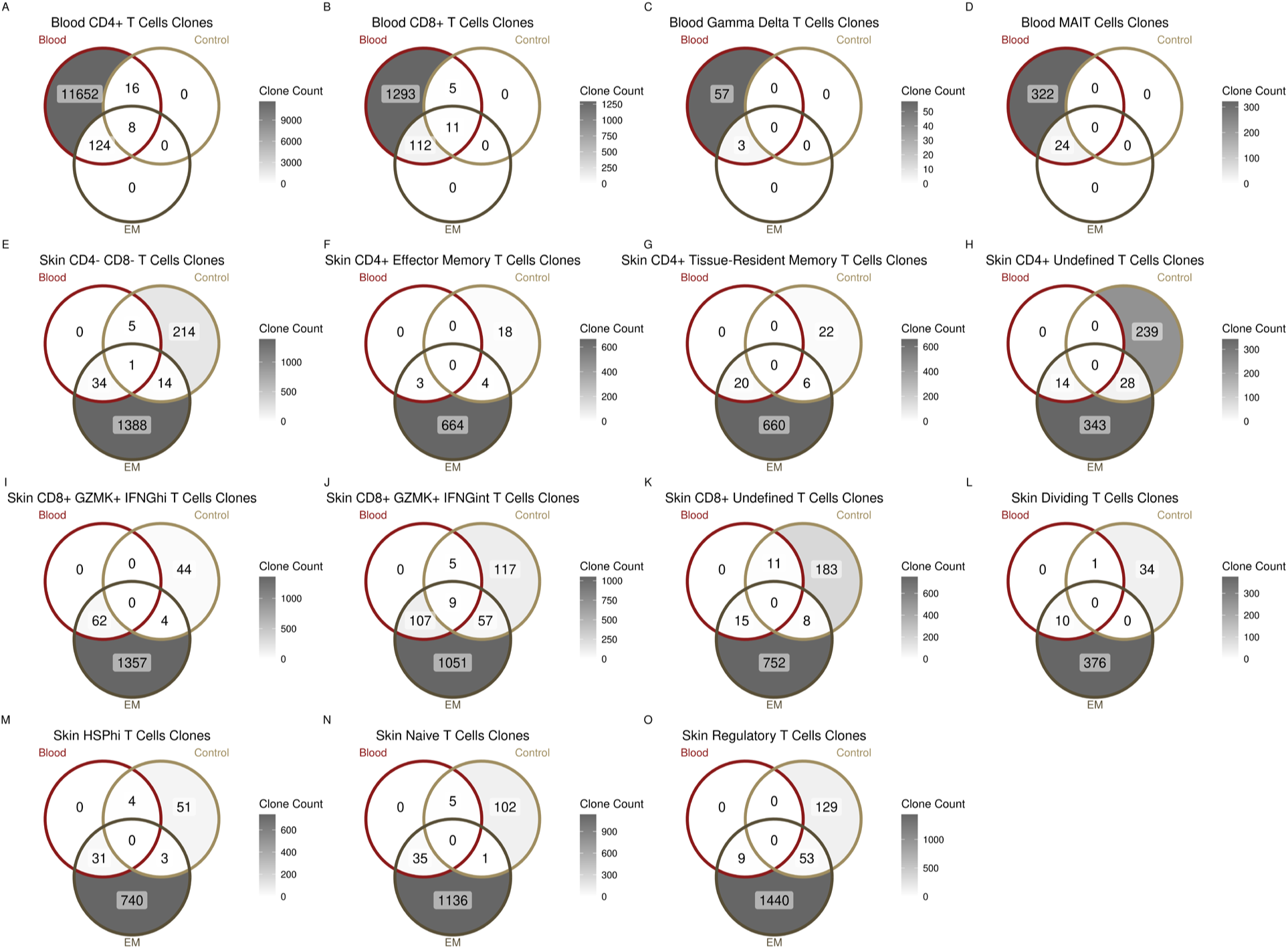
T cell clonal overlaps between the blood and the skin. **A-D)** Clonal overlaps between 4 major blood T cell subtypes (CD4^+^, CD8^+^, γδ T cells, and MAIT cells) and clonotypes within the EM and control skin samples. **E-O)** Clonotypes identified within each of the 11 defined T cell subtypes in EM and control skin samples that shared TCRs with blood samples. Numbers represent the count of unique TCRs identified and percentages represent the fraction of each segment compared to the total number of clonotypes within each respective Venn diagram.

**Supplemental Figure 8.**
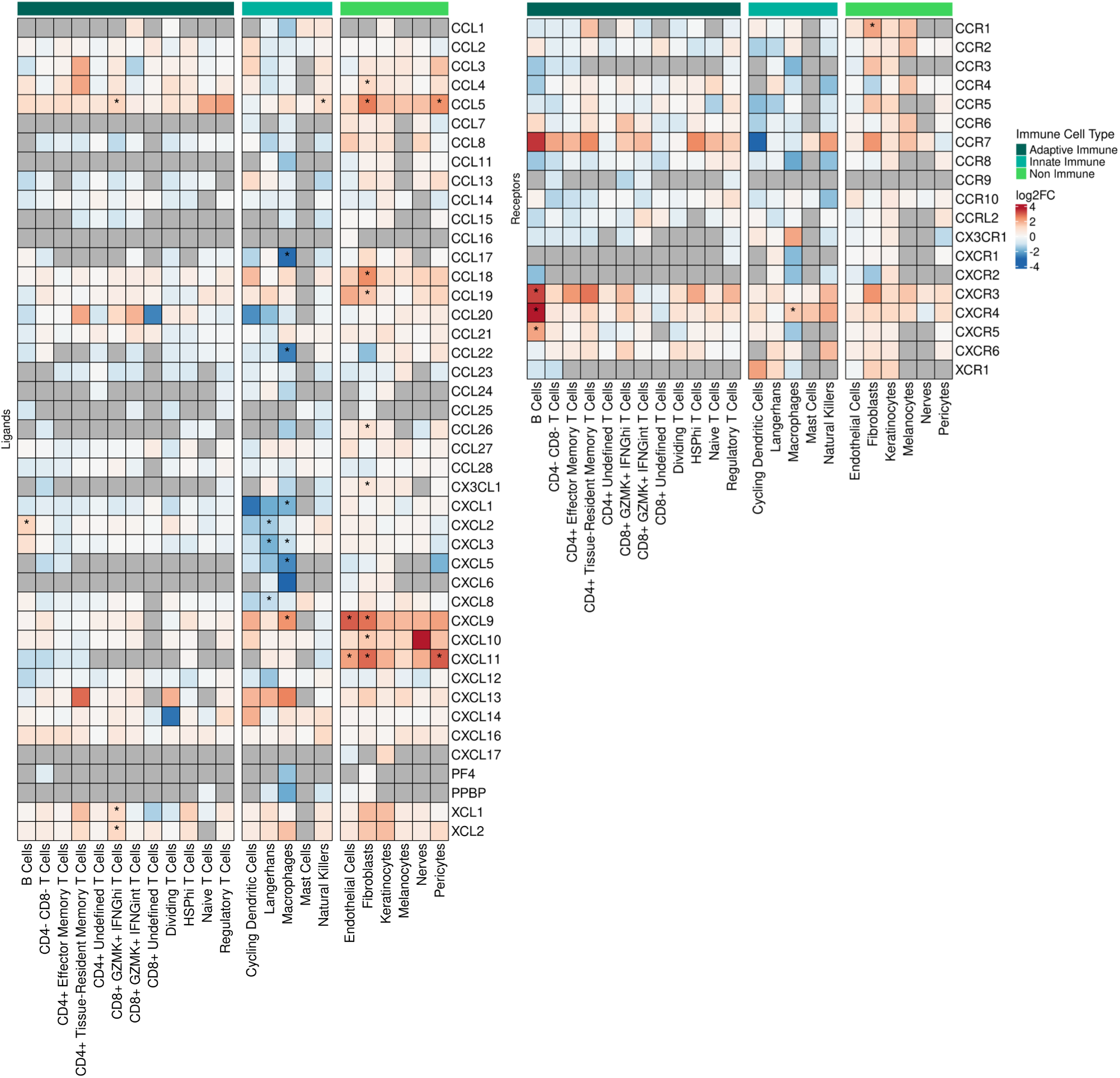
Gene expression of chemokine ligand and receptor genes in the skin. Heatmap showing the average fold change of the expression of all chemokine-related genes across the full set of skin cell types. The bar on the top indicates whether or not the cell types being plotted are immune-related (dark green for adaptive immune cell types, cyan for innate immune cell types, and bright green for non-immune cell types). Within the heatmap itself, red indicates upregulation, and blue indicates downregulation, with asterisks denoting statistically significant changes (FDR < 0.05). Gray cells indicate no expression of the relevant gene for that cell type (not applicable). Differential expression was calculated between the EM and control portions of each cell type. Note that PF4 is CXCL4 and PPBP is CXCL7.

**Supplemental Figure 9.**
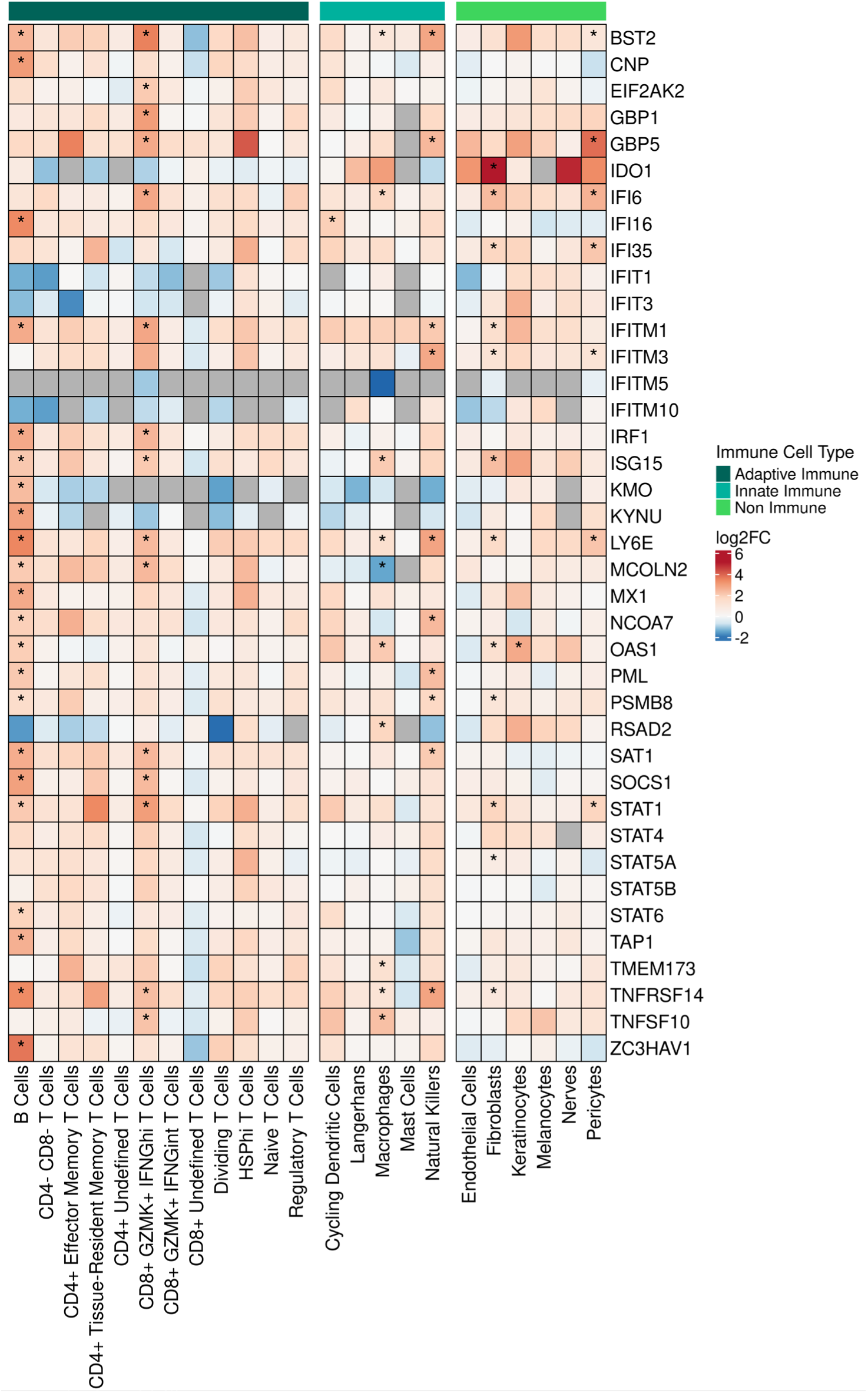
Gene expression of interferon, interferon-regulated, interleukin, and signaling-related genes in the skin. Heatmap showing the average fold change of the expression of select genes across the full set of skin cell types. The bar on the top indicates whether or not the cell types being plotted are immune-related (dark green for adaptive immune cell types, cyan for innate immune cell types, and bright green for non-immune cell types). Within the heatmap itself, red indicates upregulation, and blue indicates downregulation, with asterisks denoting statistically significant changes (FDR < 0.05). Gray cells indicate no expression of the relevant gene for that cell type (not applicable). Differential expression was calculated between the EM and control portions of each cell type.

**Supplemental Figure 10.**
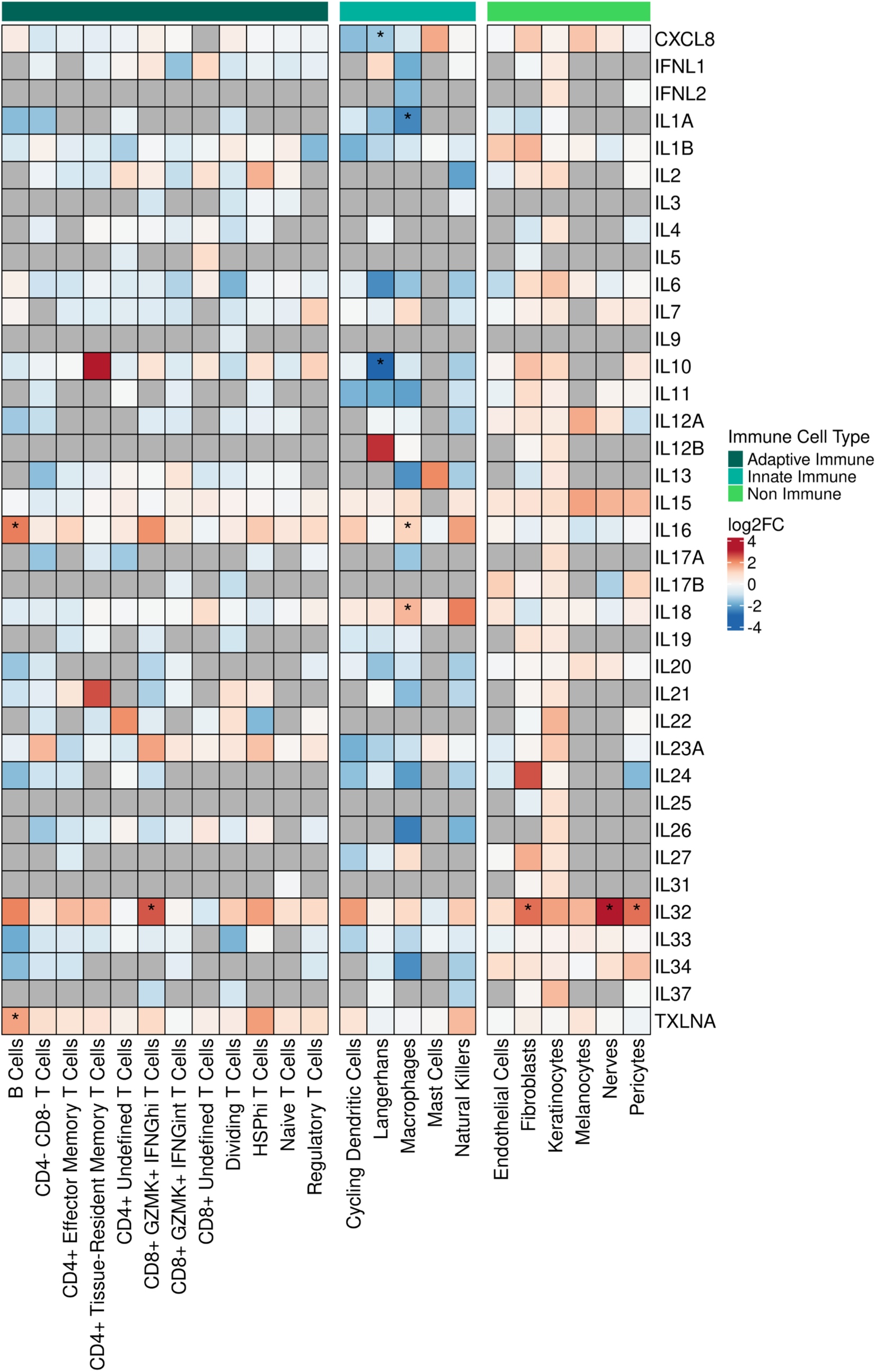
Gene expression of interleukin ligand and receptor genes in the skin. Heatmap showing the average fold change of the expression of select interleukin genes across the full set of skin cell types. The bar on the top indicates whether or not the cell types being plotted are immune-related (dark green for adaptive immune cell types, cyan for innate immune cell types, and bright green for non-immune cell types). Within the heatmap itself, red indicates upregulation, and blue indicates downregulation, with asterisks denoting statistically significant changes (FDR < 0.05). Gray cells indicate no expression of the relevant gene for that cell type (not applicable). Differential expression was calculated between the EM and control portions of each cell type. Note that CXCL8 is IL8, TXLNA is IL14, IFNL1 is IL29, and IFNL2 is IL28A.

**Supplemental Figure 11.**
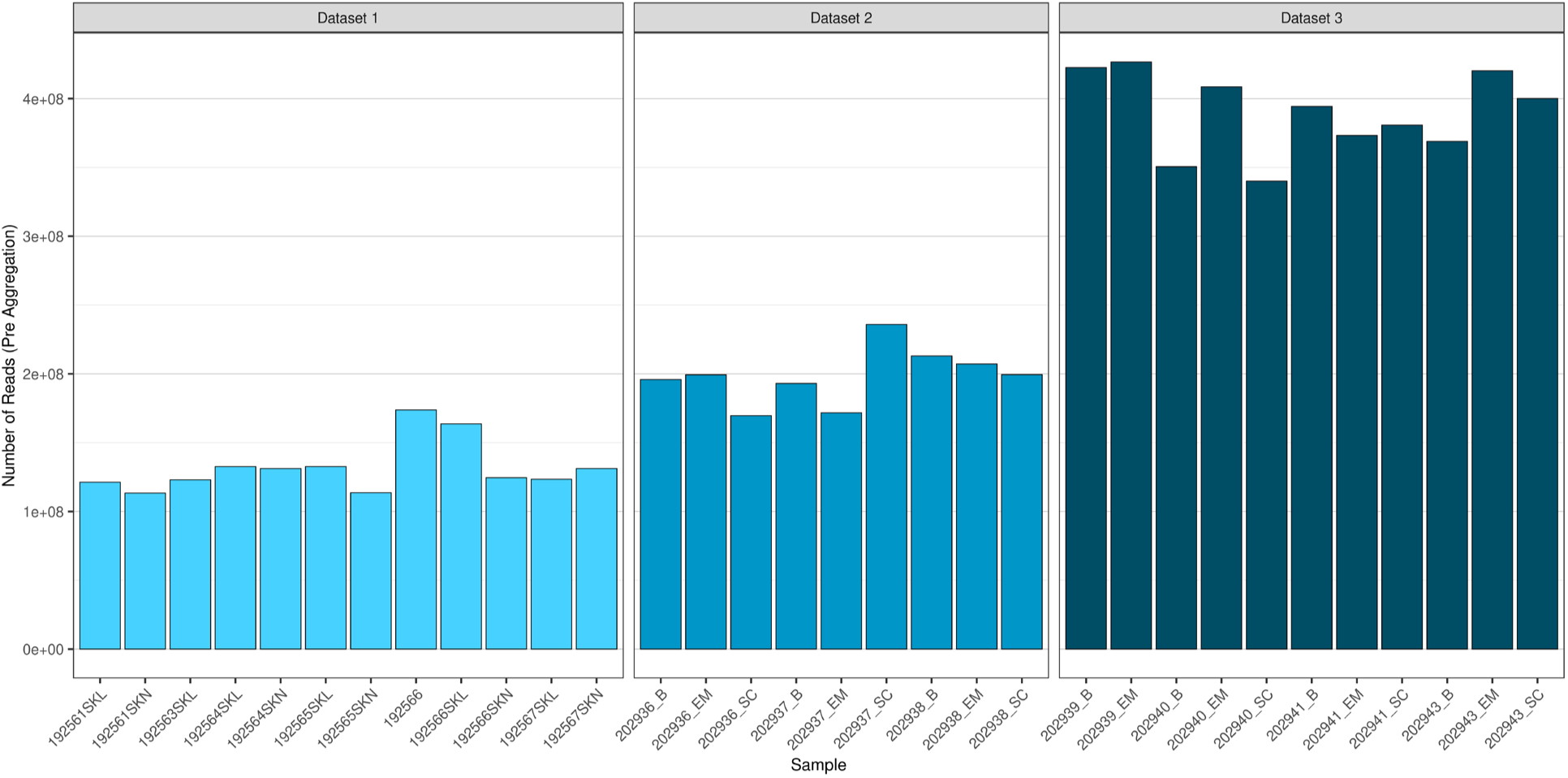
Number of reads per sample before aggregation. Bar plot showing the number of reads in the hundreds of millions for each sample (blood and skin, including control and EM) prior to aggregation, grouped and colored by dataset.

**Supplemental Figure 12.**
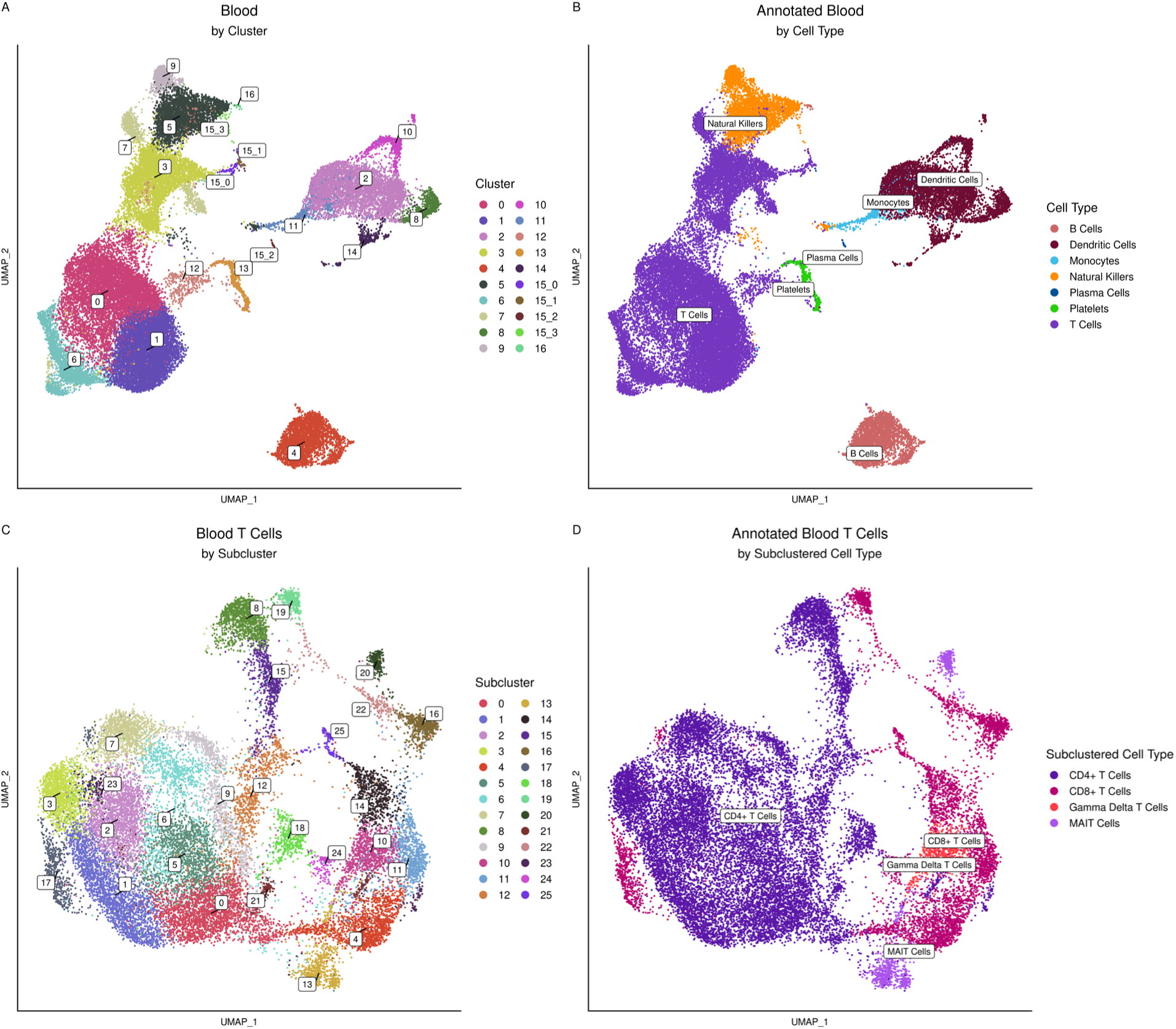
Clustering and annotation of the blood and the blood T cells. **A)** UMAP showing the Seurat clusters for the blood data. Note that cluster 15 was split into four subclusters for annotation purposes. **B)** UMAP showing the annotated blood cell types. **C)** UMAP showing the Seurat clusters for the blood T cells. **D)** UMAP showing the annotated blood T cell types.

## Notes

### Competing Interest Statement

SHK receives consulting fees from Peraton.

## References

1. Steere AC, et al. Lyme borreliosis. Nat Rev Dis Primer. 2016;2:16090.

2. Mead P. Epidemiology of Lyme Disease. Infect Dis Clin North Am. 2022;36(3):495–521.

3. Steere AC. Lyme Arthritis: A 50-Year Journey. J Infect Dis. 2024;230(Supplement_1):S1–S10.

4. Dirks J, et al. Disease-specific T cell receptors maintain pathogenic T helper cell responses in postinfectious Lyme arthritis. J Clin Invest. [published online ahead of print: July 4, 2024]. 10.1172/JCI179391.

5. Steere AC, Duray PH, Butcher EC. Spirochetal antigens and lymphoid cell surface markers in lyme synovitis. Arthritis Rheum. 1988;31(4):487–495.

6. Baarsma ME, Hovius JW. Persistent Symptoms After Lyme Disease: Clinical Characteristics, Predictors, and Classification. J Infect Dis. 2024;230(Supplement_1):S62–S69.

7. Bujak DI, Weinstein A, Dornbush RL. Clinical and neurocognitive features of the post Lyme syndrome. J Rheumatol. 1996;23(8):1392–1397.

8. Aucott JN. Posttreatment Lyme Disease Syndrome. Infect Dis Clin North Am. 2015;29(2):309– 323.

9. Marques A, et al. Transcriptome Assessment of Erythema Migrans Skin Lesions in Patients With Early Lyme Disease Reveals Predominant Interferon Signaling. J Infect Dis. 2017;217(1):158–167.

10. Lochhead RB, et al. Robust interferon signature and suppressed tissue repair gene expression in synovial tissue from patients with postinfectious, Borrelia burgdorferi-induced Lyme arthritis. Cell Microbiol. 2019;21(2):e12954.

11. Petzke MM, et al. Global Transcriptome Analysis Identifies a Diagnostic Signature for Early Disseminated Lyme Disease and Its Resolution. mBio. 2020;11(2). 10/ggp748.

12. Lochhead RB, et al. Lyme arthritis: linking infection, inflammation and autoimmunity. Nat Rev Rheumatol. 2021;17(8):449–461.

13. Glatz M, et al. Characterization of the early local immune response to Ixodes ricinus tick bites in human skin. Exp Dermatol. 2017;26(3):263–269.

14. Müllegger RR, et al. Differential Expression of Cytokine mRNA in Skin Specimens from Patients with Erythema Migrans or Acrodermatitis Chronica Atrophicans. J Invest Dermatol. 2000;115(6):1115–1123.

15. Salazar JC, et al. Coevolution of Markers of Innate and Adaptive Immunity in Skin and Peripheral Blood of Patients with Erythema Migrans. J Immunol. 2003;171(5):2660–2670.

16. Böer A, et al. Erythema migrans: a reassessment of diagnostic criteria for early cutaneous manifestations of borreliosis with particular emphasis on clonality investigations. Br J Dermatol. 2007;156(6):1263–1271.

17. Bockenstedt LK, Wooten RM, Baumgarth N. Immune Response to Borrelia: Lessons from Lyme Disease Spirochetes. Lyme Disease and Relapsing Fever Spirochetes. 2021:145–190.

18. Debes GF, McGettigan SE. Skin-Associated B Cells in Health and Inflammation. J Immunol. 2019;202(6):1659–1666.

19. Jiang R, et al. Single cell immunophenotyping of the skin lesion erythema migrans Identifies IgM memory B cells. JCI Insight. [published online ahead of print: June 1, 2021]. 10/gkct6k.

20. Zheng GXY, et al. Massively parallel digital transcriptional profiling of single cells. Nat Commun. 2017;8(1):14049.

21. Vorstandlechner V, et al. Deciphering the functional heterogeneity of skin fibroblasts using single-cell RNA sequencing. FASEB J. 2020;34(3):3677–3692.

22. Alagar Boopathy LR, et al. Mechanisms tailoring the expression of heat shock proteins to proteostasis challenges. J Biol Chem. 2022;298(5):101796.

23. Shen S, et al. Treg cell numbers and function in patients with antibiotic-refractory or antibiotic-responsive lyme arthritis. Arthritis Rheum. 2010;62(7):2127–2137.

24. Feng T, et al. Interleukin-12 Converts Foxp3+ Regulatory T Cells to Interferon–γ-Producing Foxp3+ T Cells That Inhibit Colitis. Gastroenterology. 2011;140(7):2031–2043.

25. Koenecke C, et al. IFN-γ Production by Allogeneic Foxp3+ Regulatory T Cells Is Essential for Preventing Experimental Graft-versus-Host Disease. J Immunol. 2012;189(6):2890–2896.

26. Roncarolo MG, et al. The Biology of T Regulatory Type 1 Cells and Their Therapeutic Application in Immune-Mediated Diseases. Immunity. 2018;49(6):1004–1019.

27. Fujioka S, et al. Single-cell multiomic analysis revealed the differentiation, localization, and heterogeneity of IL10+ Foxp3– follicular T cells in humans. Int Immunol. 2025;dxaf014.

28. Song Y, et al. Tr1 Cells as a Key Regulator for Maintaining Immune Homeostasis in Transplantation. Front Immunol. 2021;12. 10.3389/fimmu.2021.671579.

29. Edwards CL, et al. IL-10-producing Th1 cells possess a distinct molecular signature in malaria. J Clin Invest. 2023;133(1). 10.1172/JCI153733.

30. Chen PP, et al. Alloantigen-specific type 1 regulatory T cells suppress through CTLA-4 and PD-1 pathways and persist long-term in patients. Sci Transl Med. 2021;13(617):eabf5264.

31. Verhagen J, et al. Absence of T-regulatory cell expression and function in atopic dermatitis skin. J Allergy Clin Immunol. 2006;117(1):176–183.

32. Müllegger RR, et al. Chemokine Signatures in the Skin Disorders of Lyme Borreliosis in Europe: Predominance of CXCL9 and CXCL10 in Erythema Migrans and Acrodermatitis and CXCL13 in Lymphocytoma. Infect Immun. 2007;75(9):4621–4628.

33. Jonsson AH, et al. Granzyme K+ CD8 T cells form a core population in inflamed human tissue. Sci Transl Med. 2022;14(649):eabo0686.

34. Watanabe R, et al. Human skin is protected by four functionally and phenotypically discrete populations of resident and recirculating memory T cells. Sci Transl Med. 2015;7(279):279ra39–279ra39.

35. Henson SM, Riddell NE, Akbar AN. Properties of end-stage human T cells defined by CD45RA re-expression. Curr Opin Immunol. 2012;24(4):476–481.

36. Thome JJC, Farber DL. Emerging concepts in tissue-resident T cells: lessons from humans. Trends Immunol. 2015;36(7):428–435.

37. Knörck A, et al. Cytotoxic Efficiency of Human CD8+ T Cell Memory Subtypes. Front Immunol. 2022;13. 10.3389/fimmu.2022.838484.

38. Krummey SM, et al. CD45RB Status of CD8+ T Cell Memory Defines T Cell Receptor Affinity and Persistence. Cell Rep. 2020;30(5):1282–1291.e5.

39. Mosser DD, et al. Role of the Human Heat Shock Protein hsp70 in Protection against Stress-Induced Apoptosis. Mol Cell Biol. 1997;17(9):5317–5327.

40. Chand K, Iyer K, Mitra D. Comparative analysis of differential gene expression of HSP40 and HSP70 family isoforms during heat stress and HIV-1 infection in T-cells. Cell Stress Chaperones. 2021;26(2):403–416.

41. Figueiredo C, et al. Heat shock protein 70 (HSP70) induces cytotoxicity of T-helper cells. Blood. 2009;113(13):3008–3016.

42. Hirano N, et al. Engagement of CD83 ligand induces prolonged expansion of CD8+ T cells and preferential enrichment for antigen specificity. Blood. 2006;107(4):1528–1536.

43. Jin F, et al. The orphan nuclear receptor NR4A1 promotes FcεRI-stimulated mast cell activation and anaphylaxis by counteracting the inhibitory LKB1/AMPK axis. Allergy. 2019;74(6):1145–1156.

44. Shin H-J, et al. T-bet expression is regulated by EGR1-mediated signaling in activated T cells. Clin Immunol. 2009;131(3):385–394.

45. Suurväli J, et al. RGS16 Restricts the Pro-Inflammatory Response of Monocytes. Scand J Immunol. 2015;81(1):23–30.

46. Schmetterer KG, et al. Overexpression of PDE4A Acts as Checkpoint Inhibitor Against cAMP-Mediated Immunosuppression in vitro. Front Immunol. 2019;10:1790.

47. Filgueira L, et al. Human dendritic cells phagocytose and process Borrelia burgdorferi. J Immunol. 1996;157(7):2998–3005.

48. Shi Z, et al. The role of dermal fibroblasts in autoimmune skin diseases. Front Immunol. 2024;15. 10.3389/fimmu.2024.1379490.

49. Cavagnero KJ, Gallo RL. Essential immune functions of fibroblasts in innate host defense. Front Immunol. 2022;13. 10.3389/fimmu.2022.1058862.

50. Kersh AE, et al. CXCL9, CXCL10, and CCL19 synergistically recruit T lymphocytes to skin in lichen planus. JCI Insight. 2024;9(20). 10.1172/jci.insight.179899.

51. Duray PH. Histopathology of clinical phases of human Lyme disease. Rheum Dis Clin North Am. 1989;15(4):691–710.

52. Crawford A, et al. A Role for the Chemokine RANTES in Regulating CD8 T Cell Responses during Chronic Viral Infection. PLoS Pathog. 2011;7(7):e1002098.

53. Syed M, et al. The multifaceted role of XCL1 in health and disease. Protein Sci Publ Protein Soc. 2025;34(2):e70032.

54. Bhat MY, et al. Comprehensive network map of interferon gamma signaling. J Cell Commun Signal. 2018;12(4):745.

55. Lee J, Kim D, Min B. Tissue Resident Foxp3+ Regulatory T Cells: Sentinels and Saboteurs in Health and Disease. Front Immunol. 2022;13. 10.3389/fimmu.2022.865593.

56. Aass KR, Kastnes MH, Standal T. Molecular interactions and functions of IL-32. J Leukoc Biol. 2021;109(1):143–159.

57. Kitsou C, Fikrig E, Pal U. Tick host immunity: vector immunomodulation and acquired tick resistance. Trends Immunol. 2021;42(7):554–574.

58. Kleissl L, et al. Ticks’ tricks: immunomodulatory effects of ixodid tick saliva at the cutaneous tick-host interface. Front Immunol. 2025;16:1520665.

59. Poon MML, et al. Tissue adaptation and clonal segregation of human memory T cells in barrier sites. Nat Immunol. 2023;24(2):309–319.

60. Hamann D, et al. Phenotypic and Functional Separation of Memory and Effector Human CD8+ T Cells. J Exp Med. 1997;186(9):1407–1418.

61. Zhang F, et al. Defining inflammatory cell states in rheumatoid arthritis joint synovial tissues by integrating single-cell transcriptomics and mass cytometry. Nat Immunol. 2019;20(7):928– 942.

62. Duquette D, et al. Human Granzyme K Is a Feature of Innate T Cells in Blood, Tissues, and Tumors, Responding to Cytokines Rather than TCR Stimulation. J Immunol. 2023;211(4):633– 647.

63. Sol S, et al. Unraveling the Functional Heterogeneity of Human Skin at Single-Cell Resolution. Hematol Oncol Clin North Am. 2024;38(5):921–938.

64. Meddeb M, et al. Homogeneous Inflammatory Gene Profiles Induced in Human Dermal Fibroblasts in Response to the Three Main Species of Borrelia burgdorferi sensu lato. PLOS ONE. 2016;11(10):e0164117.

65. Ball HJ, et al. Tryptophan-Catabolizing Enzymes – Party of Three. Front Immunol. 2014;5. 10.3389/fimmu.2014.00485.

66. Opitz CA, et al. An endogenous tumour-promoting ligand of the human aryl hydrocarbon receptor. Nature. 2011;478(7368):197–203.

67. Shinde R, et al. B Cell–Intrinsic IDO1 Regulates Humoral Immunity to T Cell–Independent Antigens. J Immunol. 2015;195(5):2374–2382.

68. Strobl J, et al. Tick feeding modulates the human skin immune landscape to facilitate tick-borne pathogen transmission. J Clin Invest. 2022;132(21). 10.1172/JCI161188.

69. Berthén NC, et al. The AxBioTick study – immune gene expression signatures in human skin bitten by Borrelia-infected versus non-infected ticks. BMC Infect Dis. 2024;24(1):1422.

70. Jacobsen M, et al. Clonal Accumulation of Activated CD8+ T Cells in the Central Nervous System during the Early Phase of Neuroborreliosis. J Infect Dis. 2003;187(6):963–973.

71. Nordberg M, et al. Cytotoxic mechanisms may play a role in the local immune response in the central nervous system in neuroborreliosis. J Neuroimmunol. 2011;232(1):186–193.

72. Busch DH, et al. Detection of Borrelia burgdorferi-specific CD8+ cytotoxic T cells in patients with Lyme arthritis. J Immunol Baltim Md 1950. 1996;157(8):3534–3541.

73. Ordóñez D, et al. Cell-Mediated Cytotoxicity in Lyme Arthritis. Arthritis Rheumatol. 2023;75(5):782–793.

74. Lyme Disease (Borrelia burgdorferi) 2017 Case Definition | CDC [Internet]. 2022. https://ndc.services.cdc.gov/case-definitions/lyme-disease-2017/. Accessed September 12, 2024.

75. Song L, et al. TRUST4: immune repertoire reconstruction from bulk and single-cell RNA-seq data. Nat Methods. 2021;18(6):627–630.

76. Hao Y, et al. Integrated analysis of multimodal single-cell data. Cell. 2021;184(13):3573–3587.e29.

77. Germain P-L, et al. Doublet identification in single-cell sequencing data using *scDblFinder* [preprint]. 2022. 10.12688/f1000research.73600.2.

78. Michalik J, Niederlova V, Stepanek O. IDEIS: a tool to identify PTPRC/CD45 isoforms from single-cell transcriptomic data. Front Immunol. 2024;15:1446931.

79. Gabernet G, et al. nf-core/airrflow: An adaptive immune receptor repertoire analysis workflow employing the Immcantation framework. PLOS Comput Biol. 2024;20(7):e1012265.

80. Vander Heiden JA, et al. pRESTO: a toolkit for processing high-throughput sequencing raw reads of lymphocyte receptor repertoires. Bioinformatics. 2014;30(13):1930–1932.

81. Gupta NT, et al. Change-O: a toolkit for analyzing large-scale B cell immunoglobulin repertoire sequencing data. Bioinformatics. 2015;31(20):3356–3358.

82. Ye J, et al. IgBLAST: an immunoglobulin variable domain sequence analysis tool. Nucleic Acids Res. 2013;41(W1):W34–40.

83. Giudicelli V, Chaume D, Lefranc M-P. IMGT/GENE-DB: a comprehensive database for human and mouse immunoglobulin and T cell receptor genes. Nucleic Acids Res. 2005;33(suppl_1):D256–D261.

84. Nouri N, Kleinstein SH. A spectral clustering-based method for identifying clones from high-throughput B cell repertoire sequencing data. Bioinformatics. 2018;34(13):i341–i349.

85. Gao C-H, et al. ggVennDiagram: Intuitive Venn diagram software extended. iMeta. 2024;3(1):e177.

86. Yang Q. APackOfTheClones: Visualization of clonal expansion with circle packing. J Open Source Softw. 2024;9(103):6868.

87. Squair JW, et al. Confronting false discoveries in single-cell differential expression. Nat Commun. 2021;12(1):5692.

88. Liberzon A, et al. The Molecular Signatures Database Hallmark Gene Set Collection. Cell Syst. 2015;1(6):417–425.

89. Yu G, et al. clusterProfiler: an R Package for Comparing Biological Themes Among Gene Clusters. OMICS J Integr Biol. 2012;16(5):284–287.

90. Edgar R, Domrachev M, Lash AE. Gene Expression Omnibus: NCBI gene expression and hybridization array data repository. Nucleic Acids Res. 2002;30(1):207–210.

